# Engineered IL-10 variants elicit potent immuno-modulatory activities at therapeutic low ligand doses

**DOI:** 10.1101/2020.03.10.985069

**Authors:** C. Gorby, J. Sotolongo Bellón, S. Wilmes, W. Warda, E. Pohler, P.K. Fyfe, A. Cozzani, C. Ferrand, M.R. Walter, S. Mitra, J. Piehler, I. Moraga

**Author notes:** Corresponding authors (I.M.).

## Abstract

Interleukin-10 is a dimeric cytokine with both immune-suppressive and immune-stimulatory activities. Despite its immuno-modulatory potential, IL-10-based therapies have shown only marginal benefits in the clinic. Here we have explored whether the stability of the IL-10-receptor complex contributes to IL-10 immuno-modulatory potency. For that, we have generated an IL-10 mutant with greatly enhanced affinity for its IL-10Rβ receptor via yeast surface display. The affinity enhanced IL-10 variants recruited IL-10Rβ more efficiently into active cell surface signaling complexes than the wild-type cytokine and triggered more potent STAT1 and STAT3 activation in human monocytes and CD8 T cells. This in turn led to more robust induction of IL-10-mediated gene expression programs at a wide range of ligand concentrations in both human cell subsets. IL-10 regulated genes are involved in monocyte energy homeostasis, migration and trafficking, and genes involved in CD8 T cell exhaustion. Interestingly, at non-saturating doses, IL-10 lost key components of its gene-expression program, which may explain its lack of efficacy in clinical settings. Remarkably, our engineered IL-10 variant exhibited a more robust bioactivity profile than IL-10 wt at all the doses tested in monocytes and CD8 T cells. Moreover, CAR-modified T cells expanded with the engineered IL-10 variant displayed superior cytolytic activity than those expanded with IL-10 wt. Our study provides unique insights into how IL-10-receptor complex stability fine-tunes IL-10 biology, and opens new opportunities to revitalize failed IL-10 therapies.

## INTRODUCTION

Interleukin-10 (IL-10) is a hallmark cytokine for immune regulation that elicits potent anti-inflammatory responses. IL-10 regulates the adaptive arm of the immune response by reducing the antigen presentation potential of innate cells through decreasing their surface major histocompatibility complex (MHC) levels and co-stimulatory molecules (de Waal Malefyt et al., 1991b; Willems et al., 1994). In addition, IL-10 is a potent suppressor of the production of pro-inflammatory cytokines from a variety of cell types including monocytes, macrophages and T cells (Fiorentino et al., 1991a; Fiorentino et al., 1991b), further contributing to an anti-inflammatory environment. IL-10’s critical contribution to a healthy immune response is further highlighted by the finding that IL-10 deficient humans develop severe autoimmune diseases such as Crohn’s disease and colitis (Correa et al., 2009; Zhu et al., 2017). Despite IL-10’s relevancy for human health, the molecular bases allowing IL-10 to elicit its broad spectrum of anti-inflammatory activities are poorly understood.

Because of its potent anti-inflammatory properties recombinant IL-10 therapy was regarded as a very attractive biological approach to treat autoimmune disorders. However, despite efficacy in mouse studies (Cardoso et al., 2018; Saxena et al., 2015), IL-10 therapies failed to illicit beneficial results in the clinic, with several clinical trials showing only mild efficacy and biased responses in patients (Buruiana et al., 2010; Colombel et al., 2001). A leading hypothesis to explain the poor clinical efficacy of IL-10 is that during IL-10 therapies low levels of this cytokine reach the gastrointestinal tract, thus failing to produce an effective response (Saraiva et al., 2020). To date we have a poor understanding of how IL-10 doses influence its immuno-modulatory potential. Supporting this model, the development of strategies for a more targeted IL-10 delivery are showing enhanced clinical efficacy, although these studies are still at an early stage (Braat et al., 2006; Cardoso et al., 2018; Shigemori and Shimosato, 2017; Steidler et al., 2000). An IL-10 variant with the ability to elicit robust responses at therapeutically relevant doses would be highly desirable.

In addition to its anti-inflammatory activities, recent studies have shown that IL-10 can increase the cytotoxic function of CD8 T cells, augmenting their ability to target tumours and boosting the anti-cancer response (Oft, 2019). This seems paradoxical as IL-10 in the tumour microenvironment is linked to tumour evasion of the immune response, most likely due to IL-10’s inhibitory effects on antigen presentation (Mannino et al., 2015; Yue et al., 1997). Despite this paradox, several studies have elegantly demonstrated that IL-10 can improve production of the CD8 effector molecules granzyme B and interferon gamma both *in vitro* and *in vivo* (Emmerich et al., 2012; Mumm et al., 2011; Mumm and Oft, 2013). Currently there are several clinical trials testing the anti-tumour properties of IL-10, with already initial promising results (Naing et al., 2019). In these trials high doses of PEGylated IL-10 (Pegilodekakin) were used, which resulted in prolonged IL-10 retention in the circulation to ensure efficacy, again highlighting that effective IL-10 *in vivo* responses need high concentrations and sustained levels of IL-10.

IL-10 is a dimeric cytokine which exerts its activities by binding a surface receptor comprised of two IL-10Rα and two IL-10Rβ receptor subunits, triggering the activation of the JAK1/TYK2/STAT3/STAT1 signaling pathway and the induction of specific gene expression programs. IL-10 binds with high affinity to IL-10Rα and much lower affinity to IL-10Rβ, in the order of high micromolar/low millimolar range, (Logsdon et al., 2002). Recent studies established that sub-micromolar affinities are required to ensure efficient cytokine receptor dimerization (Richter et al., 2017; Wilmes et al., 2015).Thus, assembly of active IL-10 receptor signaling complexes in the plasma membrane is probably limited by recruitment of IL-10Rβ, making this system exquisitely sensitive to changes in either ligand concentrations and/or receptor densities in the plasma membrane. We therefore hypothesise that IL-10’s poor clinical activities result from its weak affinity for the IL-10Rβ subunit. An IL-10 variant binding IL-10Rβ with enhanced affinity has the potential to improve the therapeutic efficacy of IL-10 based therapies. To test this hypothesis, we have used the yeast surface display engineering platform to generate a new IL-10 variant which binds IL-10Rβ with 1000-fold higher affinity than IL-10 wild type (wt). Due to the dimeric nature of IL-10 wt, which makes its manipulation challenging, we have used the previously described monomeric IL-10 variant as an engineering scaffold (Josephson et al., 2000). Initially, we generated a high affinity monomeric IL-10 variant and then translated this into its natural dimeric conformation, thus obtaining four ligands to test for activity, i.e. IL-10 wild type dimer (WTD), IL-10 wild type monomer (WTM), high affinity IL-10 dimer (R5A11D) and high affinity IL-10 monomer (R5A11M). These molecules provided us with the unique opportunity to assess the contributions of IL-10 receptor binding affinity as well as IL-10 receptor complex stoichiometry to IL-10 biology. Our data showed that increasing the affinity of IL-10 for IL-10Rβ enhances IL-10’s known properties at both the molecular and cellular level. Quantitative imaging studies revealed that our high affinity variants, either in monomeric or dimeric forms, exhibited enhanced receptor heterodimerization, which in turn, resulted in a more potent activation of STAT1 and STAT3. In agreement with their improved signaling profiles, the affinity enhanced IL-10 variants induced more robust gene expression programs than the wildtype ligands in monocytes and CD8 T cells and stronger cellular responses. Accordingly, CAR-modified T cells cultured with R5A11D displayed robust cytolytic activities *in vitro* against a target leukemic cell line. Overall, our study provides novel insights into how IL-10 doses regulate its immuno-modulatory activities and show that our engineered IL-10 variants represent a clear therapeutic advantage over wild type IL-10 by eliciting more robust bioactivities at a wider range of doses.

## RESULTS

### Engineering IL-10 variants with enhanced affinity towards IL-10Rβ

IL-10 probably engages its tetrameric receptor complex in a multi-step binding process. IL-10 is recruited to the cell surface by the fast and high affinity interaction with IL-10Rα. Subsequently a second IL-10Rα subunit and two IL-10Rβ subunits, which only recognize the IL-10/IL-10Rα complex (Walter, 2014), can be recruited to initiate signalling (Figure 1A, top panel). While the exact stoichiometry of the signaling complex in the plasma membrane is still unclear, signal activation strictly requires the formation of IL-10Rα/IL-10Rβ heterodimers. A striking feature of IL-10, however, is its very poor binding affinity for IL-10Rβ (∼mM range), which we hypothesised acts as a rate-limiting step in IL-10’s biological activities. Thus, we asked whether an IL-10Rβ affinity-enhanced IL-10 variant would overcome this *in vivo* rate-limiting-step by inducing robust responses at a wide range of ligand concentrations. To address this question, we have used yeast surface display to increase the binding affinity of IL-10 for IL-10Rβ and study the signaling and activity profiles induced by these new affinity-enhanced IL-10 variants. A caveat to engineering IL-10 is its dimeric nature, which makes the correct display of this cytokine on the yeast surface challenging. We have used the monomeric IL-10 variant previously described by the Walter group (Josephson et al., 2000) as an engineering scaffold to overcome this limitation. The monomeric IL-10 was generated by extending the connecting linker between helices D and E in IL-10 by 6 peptides, consequently allowing helices E and F to fold into its own hydrophobic core to form an IL-10 monomer (Figure 1b and Sup. Figure 1). Monomeric IL-10 recruits one molecule each of IL-10Rα and IL-10Rβ to form an active signaling trimeric complex (Figure 1a, bottom panel). Although monomeric IL-10 can trigger IL-10-mediated responses, it does so with a significantly lower potency than its dimeric counterpart (Josephson et al., 2000; Logsdon et al., 2002).

**Figure 1.**
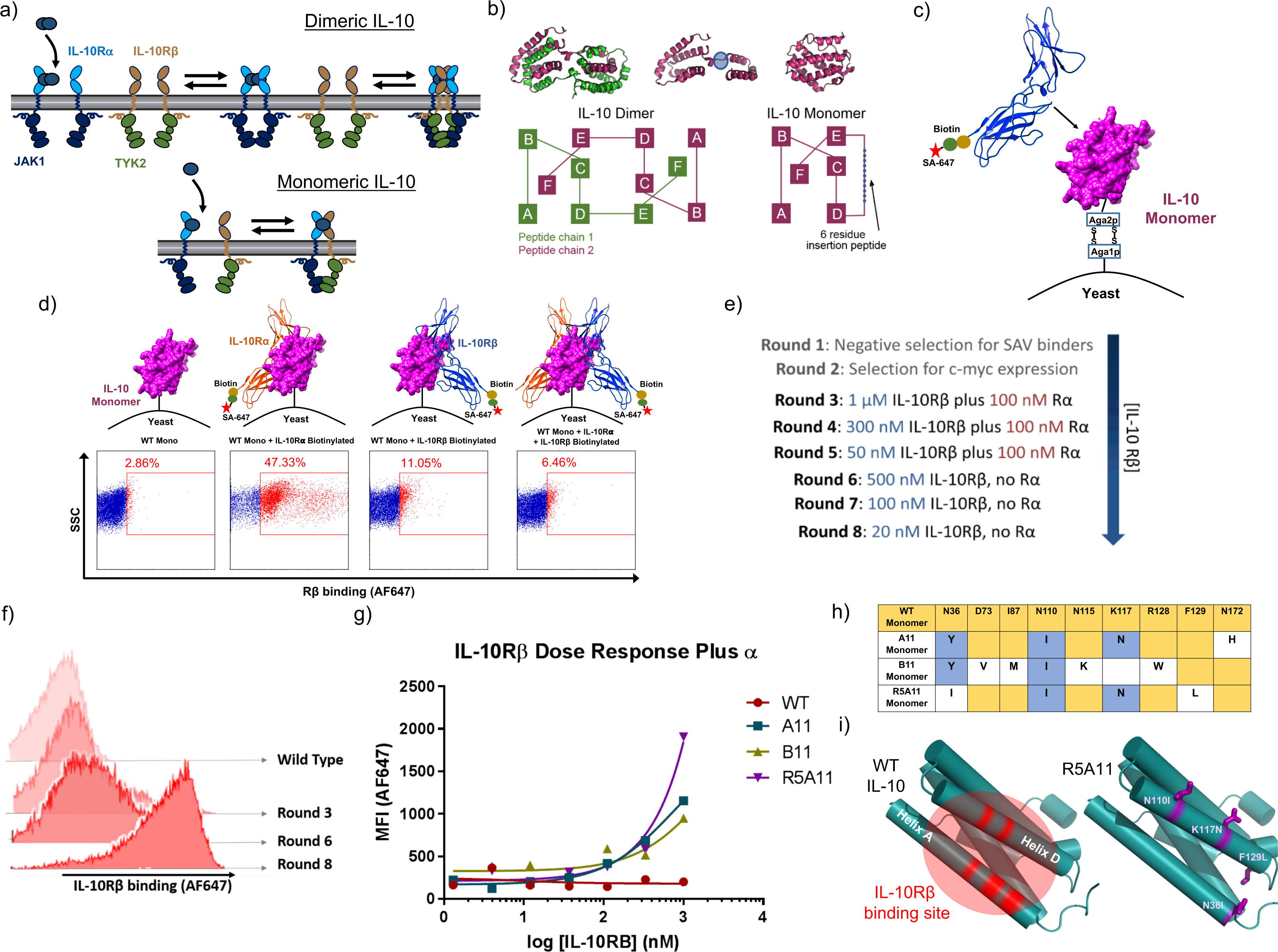
Generation of high affinity IL-10 variants by yeast surface display. **(a).** Schematic of IL-10 stepwise receptor assembly for IL-10 dimer (top panel) and IL-10 monomer (bottom panel). (**b).** Schematic of IL-10 dimer and IL-10 monomer secondary structure organisation as described by (Josephson et al., 2000; Walter, 2014). The extended linker region is highlighted in blue. (**c).** Representation of IL-10 displayed on yeast cell surface and screening using fluorescently labelled recombinant IL-10Rβ. (**d).** Yeast displayed wild type IL-10 IL-10Rα binding (panel 2) and IL-10Rβ binding in the absence (panel 3) or presence (panel 4) of IL-10Rα. Unstained control shown in panel 1. (**e).** Outline of ligand conditions used in each yeast display selection round. Selection rounds started at 1 μM IL-10Rβ with 100 nM non-biotinylated IL-10Rα and finished with 20 nM IL-10Rβ alone. (**f).** Representative histogram of IL-10Rβ binding (AF647) of yeast displayed wild type IL-10, round 3 selection, round 6 selection and round 8 selection. As the library selection proceeds the IL-10Rβ staining improves. **(g).** Dose response for IL-10Rβ binding for single clones from yeast display library. The highest IL-10Rβ concentration is 1 μM with a 1/3 serial fold dilution over 7 concentrations. Non-biotinylated IL-10Rα was added at 100 nM to improve cooperative binding. **(h).** Table for amino acid changes found in high affinity mutants. Wild type sequence is shown in yellow and is based on the amino acid sequence reported for IL-10 including leader sequence on UniProt database (P22301) with the addition of the 6 amino acid linker at position 134 allowing the formation of the monomer. Conserved changes between mutants are shown in blue. Individual mutations are shown in white. **(i).** Panel one depicts the wild type IL-10 structure with helices A and D emphasised in red as the area predicted by (Mendoza et al., 2017) to be the IL-10Rβ binding site. Panel 2 shows a cartoon representation of the IL-10 monomer with the positions of the mutations found in the high affinity variant R5A11 highlighted in purple.

First, we transfected yeast with the monomeric IL-10 construct to test whether binding to IL-10Rα and IL-10Rβ receptor subunits was preserved in the context of the yeast surface. We used biotinylated ectodomains of IL-10Rα and IL-10Rβ receptors in combination with Alexa-647 fluorescently labelled streptavidin to measure receptor binding by flow cytometry (Figure 1c). As shown in Figure 1d monomeric IL-10 retained binding to IL-10Rα confirming that it was correctly displayed on the surface of the yeast. We could not detect binding of monomeric IL-10 to IL-10Rβ in the presence or absence of IL-10Rα confirming its weak binding to this receptor subunit (Figure 1d). Without a crystal structure of IL-10 bound to IL-10Rβ to guide us in the design of a site-directed mutant library, we undertook an unbiased error-prone approach to generate IL-10 mutants with enhanced affinity for IL-10Rβ. The monomeric IL-10 variant was subject to error-prone PCR and the amplified PCR product was electroporated into the *S. cerevisiae* strain EBY100 following previously described protocols (Chao et al., 2006; Mendoza et al., 2017). Eight rounds of selection were performed where the concentration of IL-10Rβ was gradually decreased to isolate variants of IL-10 that bind IL-10Rβ with enhanced affinity (Figure 1e). Initial rounds of selection were performed with high concentrations of biotinylated IL-10Rβ in the presence of non-biotinylated IL-10Rα to stabilize the surface complex and recover low affinity binders. After round 6 the library was comprised of variants that could bind IL-10Rβ even in the absence of IL-10Rα and by round 8 the library could bind concentrations of IL-10Rβ in the low nanomolar range (Figure 1e and 1f). At this point we picked individual yeast colonies and isolated several clones (A11, B11, R5A11) that bound IL-10Rβ with enhanced affinity when compared to IL-10 wild-type (Figure 1f and 1g). When the mutations found in these variants were placed in the context of the IL-10 structure, they were found to localise in the region along helices A and D previously predicted to bind IL-10Rβ (Mendoza et al., 2017) validating our selection process (Figure 1h and 1i).

### Engineered IL-10 variants bind IL-10Rβ with nanomolar affinities

Next we recombinantly expressed the isolated IL-10 variants and characterized their biophysical properties. Importantly, the IL-10 variants behave as monomers when run in a gel filtration column confirming their monomeric nature (Sup. Figure 1a and 1b). We carried out surface plasmon resonance (SPR) studies to validate the apparent binding affinities seen in the on-yeast binding titration experiments in Figure 1g. Biotinylated IL-10Rβ was immobilised onto the chip surface and the IL-10 variants A11, B11 and R5A11 were flowed across (Sup. Figure 2a). We could not detect binding of the wild type monomeric IL-10 (WTM) at the range of doses used in this study (micromolar range), confirming the low binding affinity exhibited by IL-10 wt for IL-10Rβ (Sup. Figure 2b, panel one and 2c). The affinity matured IL-10 variants were all capable of binding IL-10Rβ with K_D_ values in the low micromolar range (Sup. Figure 2b and 2c) confirming their improved binding affinities.

**Figure 2.**
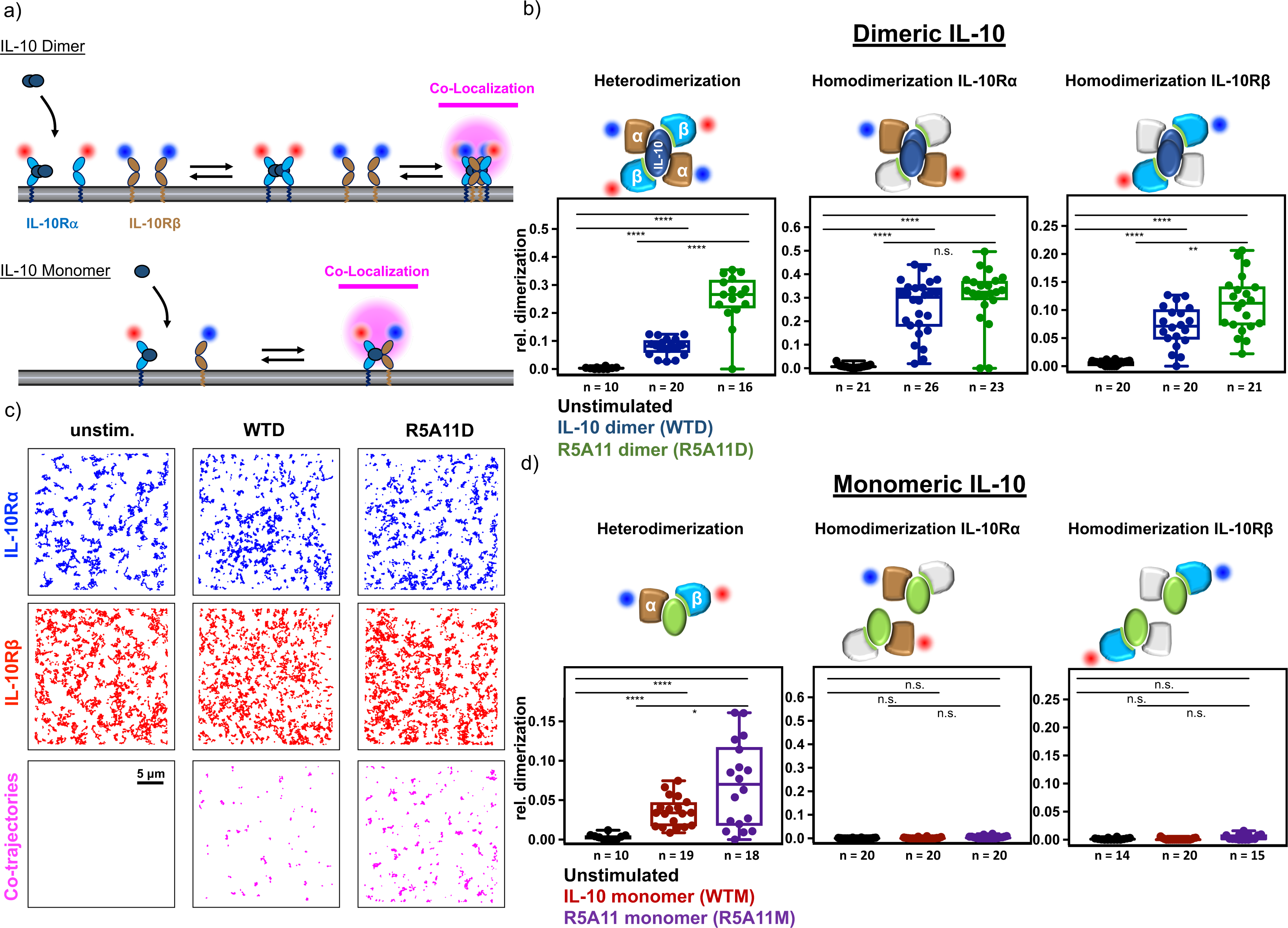
Increased IL-10Rβ binding affinity enhances IL-10 receptor dimerization. **(a).** Quantifying receptor dimerization in the plasma membrane by dual-color single molecule colocalization and co-tracking. IL-10Rα and IL-10Rβ with N-terminally fused variants of monomeric ECFP and EGFP, respectively, were labelled with nanobodies EN^AT643^ and MI^Rho11^, respectively. (**b).** Heterodimerization of IL-10R*α* and IL-10R*β* (left), and homodimerization of IL-10R*α* (center) and IL-10R*β* (right) induced by dimeric IL-10 variants quantified by co-locomotion analysis. Each data point represents a cell with the number of cells of each experiments indicated in the box plot. **c).** Trajectories of IL-10R*α* (blue), IL-10R*β* (red) and co-localized IL-10R*α*:IL-10R*β* (magenta) in the absence of IL-10 (left column) and in the presence of WTD (middle column) and R5A11D (right column), respectively. (**d).** Homo- and heterodimerization of IL-10R*α* and IL-10R*β* induced by monomeric IL-10 variants quantified by co-locomotion analysis.

IL-10 displays cooperative binding to its receptor subunits whereby its affinity for IL-10Rβ is enhanced once pre-bound to IL-10Rα (Walter, 2014). Thus, we investigated whether our mutants preserved this property. For that, we performed SPR measurements using the high affinity IL-10 variants pre-bound to soluble IL-10Rα (Sup. Figure 2d). We could not detect significant binding of the WTM/IL-10Rα complex to IL-10Rβ, highlighting again its very poor binding affinity towards IL-10Rβ (Sup. Figure 2e, panel one and 2f). All IL-10 variants exhibited enhanced binding to IL-10Rβ when complexed to IL-10Rα (in the nM range), confirming their cooperative binding and suggesting that the canonical IL-10 receptor complex binding topology has not been perturbed by the mutations introduced in our new variants (Sup. Figure 2e and 2f). These results confirm successful engineering of new IL-10 variants which engage IL-10Rβ with higher binding affinity than their wt counterpart.

### Enhanced IL-10Rβ binding affinity increases receptor heterodimerization

Thus far we had carried out the biophysical characterisation of our high affinity IL-10 variants in the monomeric conformation of the cytokine as this was necessary for the protein engineering methodologies used. In order to recapitulate the native IL-10/IL-10 receptor complex stoichiometry we recombinantly expressed our high affinity IL-10 mutant, R5A11, in the dimeric form (R5A11D) in addition to the monomeric form (R5A11M) (Sup. Figure 1a and 1b). We selected this mutant based on its higher expression yields, compared to other isolated variants. Comparisons between this and the wild type IL-10 dimer (WTD) and wild type IL-10 monomer (WTM) allowed us to examine the contributory effects of increased binding affinity as well as stoichiometry on IL-10’s molecular and cellular activities.

In order to test how increasing the binding affinity to IL-10Rβ supported receptor assembly at the plasma membrane of live cells, we probed diffusion and interaction of both receptor chains by dual colour total internal reflection fluorescence (TIRF) microscopy. To this end, we expressed in HeLa cells IL-10Rα and IL-10Rβ N-terminally tagged with engineered variants of non-fluorescent (Y67F) mEGFP. The tags were designed to specifically recognise either one of two different anti-GFP nanobodies (Kirchhofer et al., 2010). These nanobodies (NBs) were conjugated to photostable organic fluorophores ATTO Rho11 and ATTO 643 suitable for simultaneous dual-colour single molecule tracking of IL-10Rα and IL-10Rβ in the plasma membrane of live cells as shown previously in other cytokine receptor systems (Martinez-Fabregas et al., 2019; Moraga et al., 2015a; Wilmes et al., 2020) (Figure 2a and Sup. Figure 3a).

**Figure 3.**
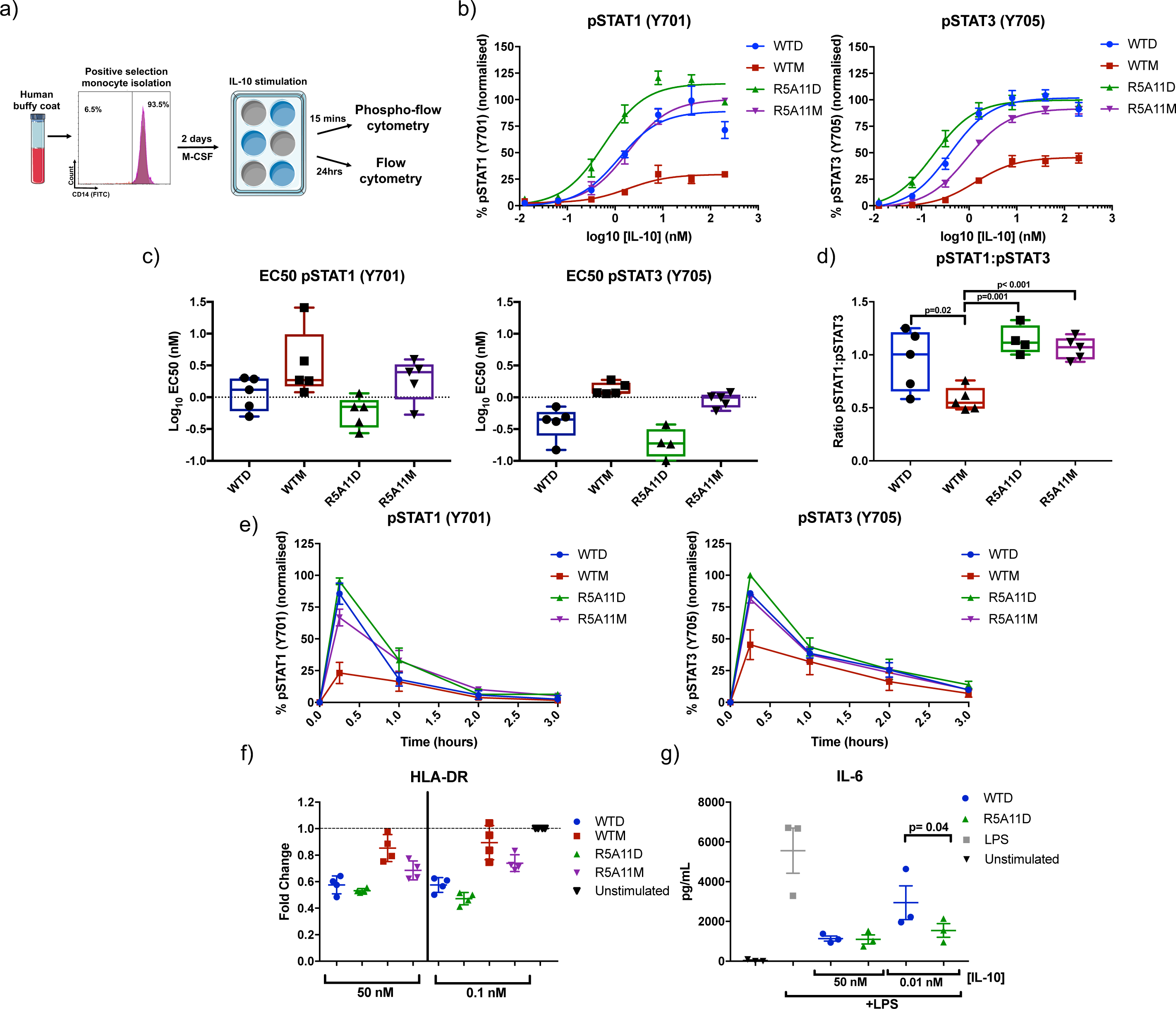
High affinity variants improve signaling capabilities of IL-10 in monocytes. **(a).** Monocytes were isolated from human buffy coat samples by CD14 positive MACS selection. Cells were rested in monocyte colony sitmualting factor (M-CSF) containing media for 2 days. Cells were then stimulated with IL-10 for 15 mins for phospho-flow cytometry analysis or 24 hours for analysis of HLA-DR levels. **(b).** Dose response of pSTAT1 and pSTAT3 in IL-10 treated monocytes. Cells were stimulated with IL-10 wild type and high affinity variants for 15 minutes. Activation of STAT1 and STAT3 was analysed by phospho-flow cytometry. Sigmoidal curves were fitted with GraphPad Prism software. Data shown is the mean of five biological replicates with error bars depicting standard error of the mean. Each biological replicate is normalised by assigning the highest MFI value of the top concentration as 100% and the lowest MFI value of an untreated control as 0%. **(c).** Log_10_ EC50 values for pSTAT1 and pSTAT3 from dose response curves in (b). Each point represents one biological replicate with line at the mean and error bars show the mix to max of all points. **(d).** Ratio of pSTAT1 to pSTAT3 in IL-10 stimulated monocytes. Ratio was calculated by taking the percentage activation of pSTAT3 and pSTAT1 at 40 nM for five biological replicates and dividing pSTAT1 by pSTAT3 values. Each point represents one biological replicate with line at the mean and error bars show the min to max of all points. P value calculated by two-tailed paired t-test. **(e).** Kinetics of pSTAT3 and pSTAT1 induced by IL-10. Monocytes were stimulated with IL-10 for the indicated time periods before fixation. Data shown is the mean of four biological replicates with error bars depicting standard error of the mean. Each biological replicate is normalised by assigning the highest MFI value at 15 mins as 100% and the lowest MFI value of an untreated control as 0%. **(f).** Measurement of HLA-DR cell surface expression in monocytes after 24 hours IL-10 treatment. Each point represents one biological replicate (n=4) and error bars indicate the standard deviation. Fold change is calculated for each biological replicate by dividing the MFI of the treated samples by a non-IL-10 treated control (unstimulated). **(g).** Monocytes were stimulated with LPS for 8 hours in the presence of IL-10. Each point represents one biological replicate (n=3) and error bars indicate the standard error of the mean. P value calculated by two-tailed ratio paired t-test.

After cell surface labelling we found both receptor subunits randomly diffusing in the plasma membrane (Figure 2b). Dimerization of receptor subunits was quantified by co-tracking analysis. Receptors were considered as dimerized if two individual particles in both spectral channels persistently co-localized for *≥*10 consecutive steps (∼320 ms) in a proximity of 150 nm. These co-localization/co-tracking thresholds allowed reliable elimination of density-dependent random co-localizations (Wilmes et al., 2020). In the absence of IL-10, we did not observe heterodimerization of IL-10Rα and IL-10Rβ (Figure 2b and 2c). Stimulation with saturating concentrations of IL-10 WTD significantly dimerized IL-10Rα and IL-10Rβ. Strikingly, IL-10 R5A11D induced a substantially higher level of receptor heterodimers (Figure 2b and 2c). This finding was also confirmed for the monomeric versions of both wild type and high affinity IL-10 variants although at lower levels than seen for the dimeric versions (Figure 2d). This observation is in line with the 50% reduced probability to observe heterodimers expected for the monomeric vs. the dimeric ligand. Ligand stimulation led to a significant decrease of diffusion mobility, particularly for IL-10Rα (Sup. Figure 3b), which is in line with previous reports on cytokine receptor dimerization (Moraga et al., 2015c; Richter et al., 2017; Wilmes et al., 2015).

We also probed homodimerization of IL-10Rα and IL-10Rβ, respectively. To this end, we stochastically labelled either of the receptor chains with both dyes (Sup. Figure 3a) taking into account that only half of the dimers would be labelled with different dyes and thus would be picked up by co-tracking analysis. Stimulation with the dimeric IL-10 induced strong homodimerization of IL-10Rα with no difference between both cytokine variants, as the IL-10Rα binding interface was unaltered in R5A11 (Figure 2c). Instead, homodimerization of IL-10Rβ was significantly increased for the engineered variant R5A11D compared to WTD. For the monomeric IL-10 variants, all homodimerization experiments failed to induce receptor homodimers, in agreement with the monomeric nature of the ligands (Figure 2d) (Josephson et al., 2000). Taken together, these results confirmed that, compared to WT IL-10, increased recruitment of IL-10Rβ into the signaling complex at the plasma membrane was achieved for the engineered R5A11 variants.

### IL-10 variants exhibit enhanced signaling activities in human primary monocytes

IL-10 inhibits inflammatory processes by modulating the activities of different innate cells including monocytes. We next performed a battery of signaling and activity assays in human monocytes to investigate the anti-inflammatory potential of our engineered variants. Monocytes (CD14^+^cells) were isolated from human buffy coats and rested for two days before stimulation with wild type IL-10 and high affinity monomer and dimers (Figure 3a). Levels of STAT1 and STAT3 phosphorylation upon ligand stimulation were measured by flow cytometry as these two transcription factors represent the major signaling pathway engaged by IL-10 (Finbloom and Winestock, 1995; Wehinger et al., 1996). At saturating concentrations R5A11D and WTD activated comparable STAT1 and STAT3 levels (Figure 3b). However, R5A11D showed enhanced phosphorylation of both STAT1 and STAT3 at sub-saturating concentrations, which translated into a decrease in EC_50_ values compared to WTD (Figure 3b and 3c). WTM showed a poor activation of STAT1 and STAT3 with amplitudes of activation reaching less than fifty percent of those elicited by WTD (Figure 3b). While WTD, R5A11D and R5A11M showed a similar pSTAT1 to pSTAT3 ratio, WTM’s clear bias towards pSTAT3 (Figure 3d), agrees with previous observations from our laboratory describing biased signaling by short-lived cytokine-receptor complexes (Martinez-Fabregas et al., 2019). R5A11M induced activation of both STAT1 and STAT3 to levels comparable to those induced by the dimeric cytokines at saturating doses, suggesting that the defective signaling elicited by WTM results from its weak IL-10Rβ binding affinity (Figure 3b and 3c). Time-dependent analysis showed that the signaling profiles of the variants were not caused by differences in the signaling kinetics. The four IL-10 ligands triggered comparable signaling kinetics in human monocytes (Figure 3e and Sup. Figure 4), confirming that their different signaling profiles result from their different binding affinities to IL-10Rβ.

**Figure 4.**
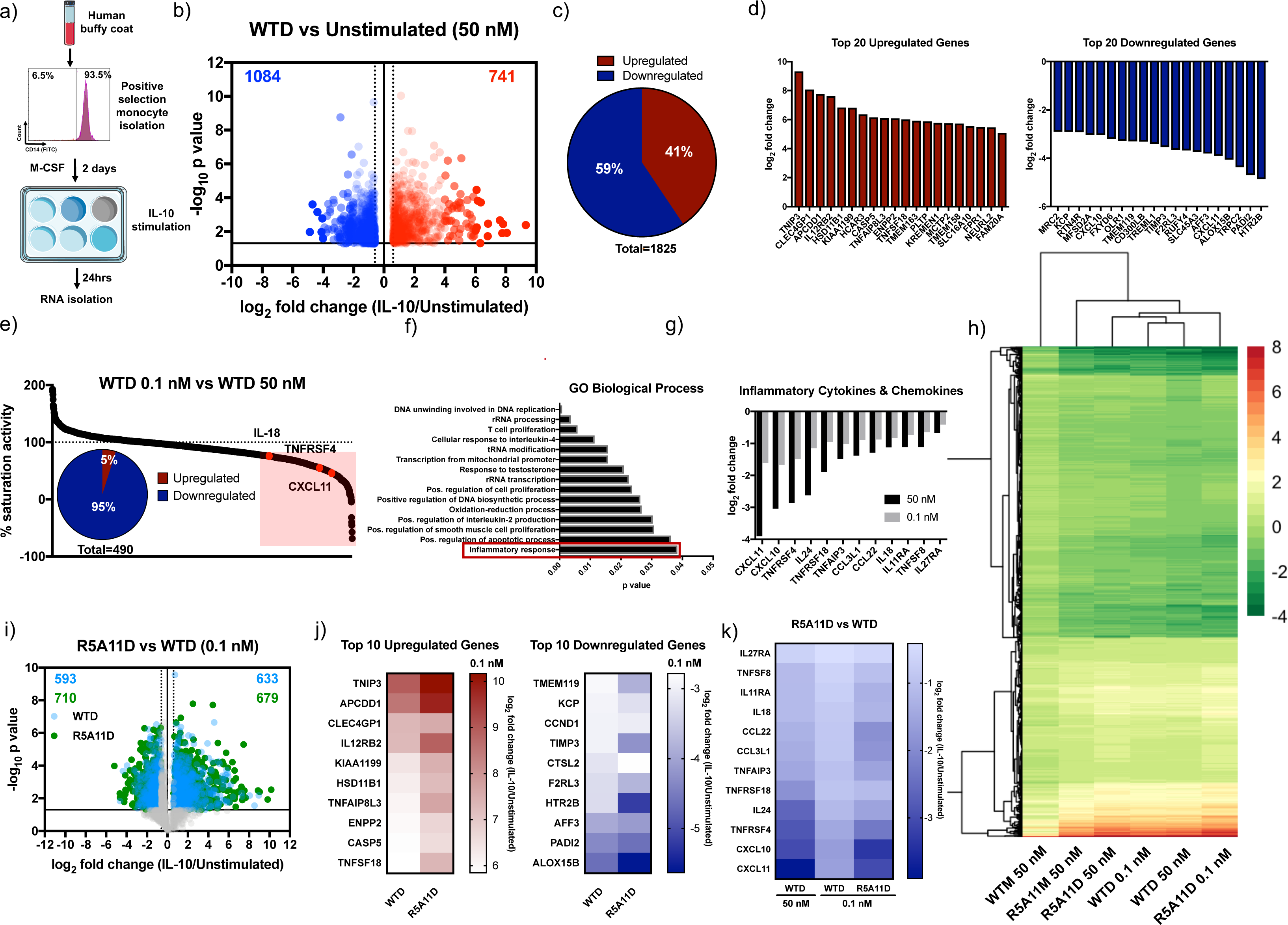
Characterisation of transcriptional activity induced by IL-10 and high affinity variants in human monocytes. **(a).** Schematic of monocyte stimulation. CD14 positive cells were isolated from three human buffy coats by MACS and rested in M-CSF containing media for two days before twenty-four hours stimulation with IL-10 wild type and high affinity variants. **(b).** Volcano plot of monocyte genes significantly upregulated by IL-10 wild type dimer ≧ 0.6 log_2_ fold change (red) and significantly downregulated ≦ −0.6 log_2_ fold change compared to non-IL-10 stimulated cells. Fold change was calculated by dividing WTD 50 nM by unstimulated values for each donor. The average fold change was calculated and the log_2_ of this value is plotted. P values ≦ 0.05 were calculated by two-tailed unpaired t-test of the log_2_ fold change of WTD 50 nM/unstimulated genes for each donor. Genes which were not significantly changed or were ≦ 0.6 ≧ −0.6 log_2_ fold change were excluded. **(c).** Proportion of monocyte genes significantly regulated by WTD 50 nM ≧ 0.6 or ≦ −0.6 log_2_ fold change compared to unstimulated cells. **(d).** Log_2_ fold change for the top 20 protein coding genes significantly up (red) and down (blue) regulated by WTD 50 nM in monocytes. **(e).** Percentage activity of low dose WTD compared to high dose WTD. The log_2_ fold change of WTD 0.1 nM was divided by WTD 50 nM and multiplied by 100. Genes which showed ≦ 75% of high dose activity (490 genes) are highlighted in red. Insert shows the percentage of these genes which were up or downregulated by IL-10. **(f).** Gene ontology biological processes analysis for the 490 genes which were sensitive to changes in WTD concentration. **(g).** Log_2_ fold change for inflammatory cytokine and chemokine genes for WTD 50 nM and 0.1 nM. **(h).** Heatmap of genes significantly up or down regulated by WTD 50 nM ≧ 0.6 or ≦ −0.6 log_2_ fold change and their corresponding log_2_ fold change by R5A11D 50 nM, WTD 0.1 nM, R5A11D 0.1 nM, R5A11M 50 nM and WTM 50 nM compared to unstimulated control cells. Values are the mean of three donors. Heat map cluster analysis generated in R studio. **(i).** Volcano plot of genes regulated by WTD (blue) and R5A11D (green) at 0.1 nM concentration each. Only genes which had already been showed to be significantly up or down regulated by WTD 50 nM are plotted. **(j).** Heatmap of the top 10 up and down regulated genes by WTD 0.1 nM compared to R5A11D 0.1 nM. **(k).** Heatmap of inflammatory cytokine and chemokine genes regulated by WTD at 50 nM and 0.1 nM and R5A11D 0.1 nM.

IL-10 exerts its anti-inflammatory properties by inhibiting antigen presentation in innate cells such as monocytes and dendritic cells (Mittal and Roche, 2015). Thus, we next studied whether IL-10 binding affinity to IL-10Rβ influences its ability to decrease HLA-DR expression in human primary monocytes. WTD and R5A11D reduced the levels of HLA-DR surface levels to a similar extent (50%) at saturating doses, in agreement with their comparable signaling profiles (Figure 3f). At sub-saturating doses however, R5A11D induced a stronger downregulation of HLA-DR expression (Figure 3f). WTM induced a mild reduction of HLA-DR surface levels (20%) paralleling its poor signaling potency (Figure 3f). R5A11M induced only a 30% reduction of the surface HLA-DR levels, despite activating STAT1/STAT3 to a very similar extent as the dimeric ligands (Figure 3f), suggesting an additional dimer-dependent mechanism by which IL-10 regulates HLA-DR expression. We next investigated how IL-10Rβ binding affinity correlates with IL-10’s ability to inhibit pro-inflammatory cytokine production by monocytes. For this, we measured levels of IL-6 secreted by monocytes upon LPS stimulation in the presence of the indicated doses of WTD and R5A11D (Fig 3g). At saturating concentrations WTD and R5A11D effectively inhibited IL-6 secretion to a similar extent (Figure 3g). However, at sub-saturating doses R5A11D again showed a marked improvement over WTD (Figure 3g). Together, our data highlight that IL-10 variants exhibiting enhanced binding towards IL-10Rβ gain a functional advantage at sub-saturating doses, such as those that would be attained during therapeutic interventions.

### Increased IL-10Rβ affinity enhances transcriptional activity of IL-10 in monocytes

Our initial studies in monocytes were focused on two classical markers regulated by IL-10, i.e. HLA-DR levels and IL-6 expression. To gain a broader understanding of how our variants regulate human monocyte activities, we performed detailed transcriptional analysis of human monocytes stimulated with the different IL-10 ligands for 24 hrs. Monocytes were isolated and treated as in Figure 4a. WTD treatment elicited a strong transcriptional regulation in human monocytes inducing the upregulation of 741 genes and downregulation of 1084 genes (Figure 4b and 4c). Highly upregulated and downregulated genes are shown in Figure 4d. KEGG pathway analysis showed a significant number of genes regulated by IL-10 treatment involved in pyruvate metabolism (Sup. Figure 5a). We also noted a number of other key metabolic genes significantly changed upon IL-10 stimulation, a selection of which are shown in Sup. Figure 5b. WTD treatment regulated expression of hexokinase-2 and hexokinase-3, key enzymes in glycolysis. Genes associated with acyl-CoA synthesis, ACSS2, ACSL4, ACSL1, were also significantly upregulated highlighting a potential regulation of lipid biosynthesis by IL-10 (Sup. Figure 5b). In addition to metabolic-related genes, WTD treatment regulated expression of cytokines, chemokines and their receptors (Sup. Figure 5b). For instance, cytokines receptors such as IL-12Rβ2, IL-21R*α* and IL-4R*α* were upregulated while cytokines such as IL-8, IL-18 and IL-24 were downregulated (Sup. Figure 5b). Expression of CXCL1, CCL22, CCL24, CCL18, CXCL10 and CXCL11 chemokines were also modulated by IL-10 treatment and can be expected to contribute to the production of an anti-inflammatory environment. We also observed the regulation of a wide variety of genes encoding cell surface proteins by WTD treatment, including CD93 – a receptor critical for monocyte phagocytosis and CD44 and CD9 – markers involved in cell surface adhesion (Sup. Figure 5b); and an inhibition of type I IFN gene signature, in agreement with previous studies (Dallagi et al., 2015; Ito et al., 1999) (Sup. Figure 5b). Overall, our transcriptome study revealed a broad regulation of monocyte biology by IL-10, that includes fine-tuning energy homeostasis, migration and trafficking. Remarkably, 73% of genes regulated by WTD at saturating doses were induced to a similar extent when sub-saturating doses of WTD were used, highlighting the robustness of the IL-10 response (Figure 4e). However, of the 27% of genes differentially regulated by WTD at the two doses investigated (0.1 nM and 50 nM), 95% of those correspond to genes downregulated by IL-10 treatment and include critical pro-inflammatory chemokines and cytokines (Figure 4e and 4f). A list of differentially expressed genes is provided in Figure 4g. Our data show that at low doses IL-10 looses the ability to block expression of key cytokines and chemokines that critically contribute to enhance the inflammatory response.

**Figure 5.**
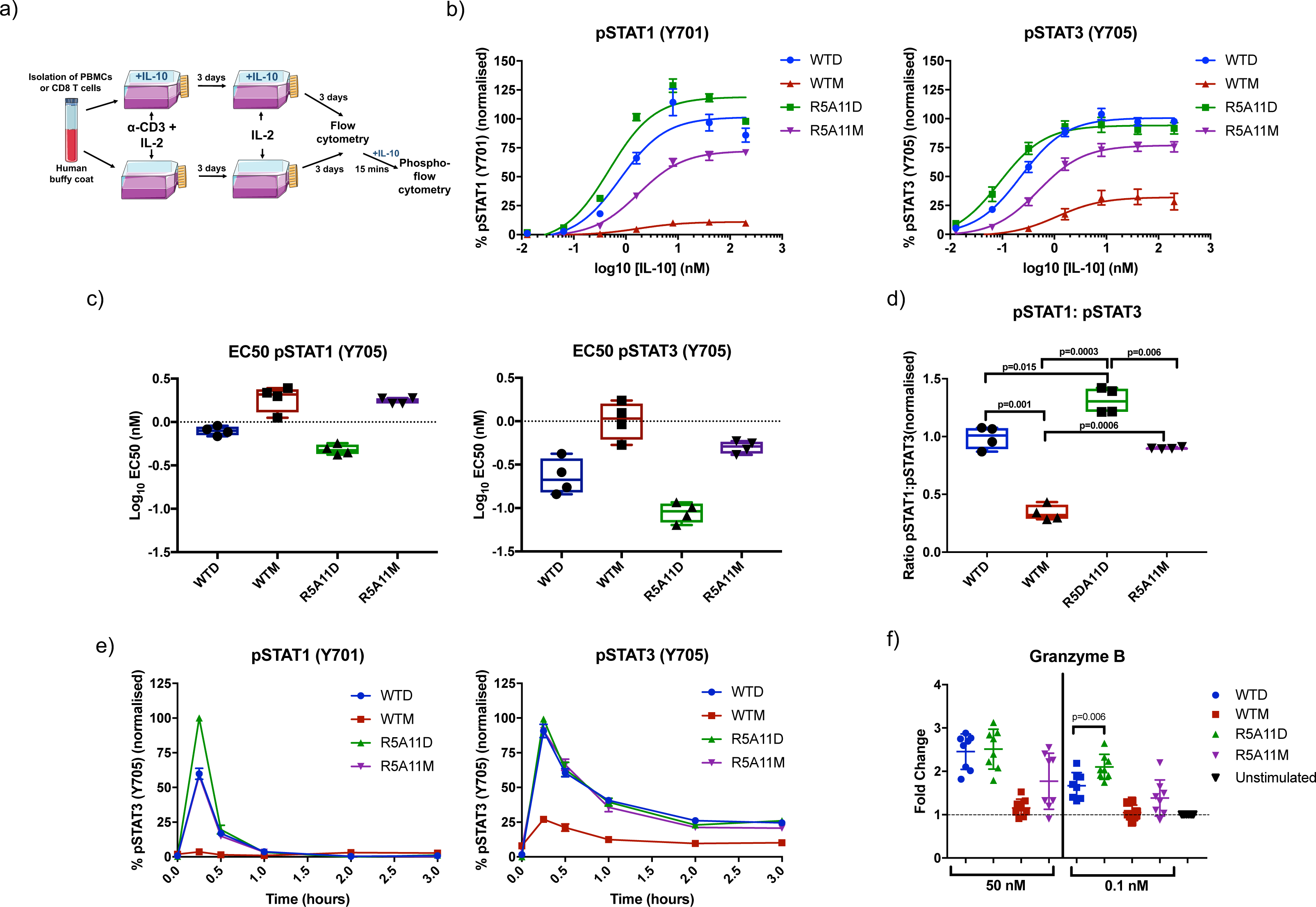
High affinity variants improve signaling capabilities of IL-10 in human CD8 T cells. **(a).** PBMCs were isolated from human buffy coat samples and CD8 cells were purified by CD8 positive MACS selection. PBMCs or purified CD8 cells were activated for three days using soluble anti-CD3 (100 ng/mL) (PBMCs) or anti-CD3/anti-CD28 beads (CD8 cells) with IL-2 (20 ng/mL) in the presence or absence of IL-10. On day 3 activation media was removed and the cell populations were placed in media containing IL-2 plus/minus IL-10 for a further 2-3 days before analysis. **(b).** Dose response of pSTAT3 and pSTAT1 in activated CD8 cells in a PBMC population (activated in the absence of IL-10). Cells were placed in media with no IL-2 overnight before stimulation with IL-10 wild type and mutants for 15 minutes. Data shown is the mean of four biological replicated with error bars depicting standard error of the mean. Each biological replicate is normalised by assigning the highest MFI value of the top concentration as 100% and the lowest MFI value of an untreated control as 0%. **(c).** Log_10_ EC50 values for pSTAT3 and pSTAT1 from dose response curves in (b). Each point represents one biological replicate with line at the mean and bars represent the mix to max of all points. **(d).** Ratio of pSTAT1 to pSTAT3 in IL-10 stimulated CD8 cells in a PBMC population. Ratio was calculated by taking the percentage activation of pSTAT3 and pSTAT1 at 40 nM for four biological replicates and dividing pSTAT1 by pSTAT3 values. Each point represents one biological replicate with line at the mean and error bars denote mix to max of all points. P value calculated by paired t tests. **(e).** Kinetics of pSTAT3 and pSTAT1 induced by IL-10. Non-activated CD8 cells in a PBMC population were stimulated with IL-10 for the indicated time periods before fixation. Data shown is the mean of three biological replicates with error bars depicting standard error of the mean. Each biological replicate is normalised by assigning the highest MFI value at 15 mins as 100% and the lowest MFI value of an untreated control as 0%. **(f).** Granzyme B protein in activated CD8 T cells in the presence of IL-10. CD8 T cells in a PBMC population were grown and stimulated as shown in (a). Cells were then fixed and permeabilised and granzyme B protein was quantified by flow cytometry. Fold change was calculated by normalising to a non-IL-10 treated control for each donor. Each point represents one biological replicate (n=8) and error bars indicate the standard deviation.

Next we studied how the engineered IL-10 variants regulated gene expression programs in monocytes. WTM induced a very poor transcriptional response, in line with its weak signal activation profile (Figure 4h and Sup. Figure 5c). Interestingly R5A11M triggered a more potent transcriptional response when compared to WTM but failed to reach the same potency induced by the dimeric ligands (Figure 4h and Sup. Figure 5c). A direct comparison between WTD and R5A11M showed that the latter monomeric ligand exhibited a diminished expression of 39% of genes regulated by WTD (Sup. Figure 5d). This is in contrast to its ability to activate STAT1 and STAT3 to levels comparable to those induced by the dimeric ligands, suggesting that STAT activation does not directly correlate with transcriptional activity in the IL-10 system. In agreement with our signaling studies, R5A11D induced a more robust gene expression profile at sub-saturating doses when compared to WTD (Figure 4h). R5A11D enhanced the expression of 18% of genes versus WTD at 0.1 nM, with only 6% of genes showing increased activity by WTD over R5A11D (Figure 4h and 4i, Sup. Figure 5e). Figure 4j shows that of the top 10 IL-10 regulated genes, the majority of them displayed enhanced sensitivity to R5A11D treatment. This pattern holds true when genes are grouped by families, i.e. cytokines & chemokines, CD markers and MAPK signaling (Sup. Figure 5c). Importantly, key pro-inflammatory cytokines, which were not regulated by WTD at low doses, are still regulated by R5A11D (Figure 4k). Overall, our transcriptional data shows that IL-10 regulates monocyte biology at different levels and that R5A11D, by exhibiting enhanced affinity towards IL-10R*β*, elicits more robust responses at a broader range of ligand concentrations, holding the potential to rescue IL-10 based therapies targeting inflammatory disorders.

### IL-10 variants exhibit enhanced signaling activities in human primary CD8 T cells

In addition to its potent anti-inflammatory effects IL-10 also stimulates cytotoxic CD8 T cells under specific circumstances, enhancing production of effector molecules and increasing their cytotoxic activity (Oft, 2014). We next investigated whether the enhanced activities exhibited by our affinity-matured variants in monocytes would translate into CD8 T cells. Human primary CD8 T cells were grown and activated as shown in Figure 5a and STAT1/STAT3 activation levels in response to the indicated concentrations of IL-10 variants were measured by flow cytometry (Figure 5b). WTD and R5A11D induced very similar STAT phosphorylation levels at saturating doses, but R5A11D showed a decreased EC_50_ value and stronger signaling at sub-saturating doses (Figure 5b-d), agreeing with our results in monocytes. R5A11D showed a more potent activation of STAT1 over STAT3 which we did not observed in monocytes, suggesting that long-lived IL-10 receptor complexes gain an advantage activating STAT1 in CD8 T cells. WTM exhibited weak activation of STAT1 and STAT3, inducing less than 25% of the activation amplitudes elicited by the dimeric molecules and exhibited a biased STAT3 activation (Figure 5b-d). In contrast to what we observed in monocytes, R5A11M also elicited a STAT3 biased response activating STAT3 to 80% of the levels induced by the dimeric molecules and STAT1 to 60% of the levels induced by the dimeric molecules (Figure 5b-d), suggesting that fundamental differences between monocytes and CD8 T cells impact signaling downstream of the IL-10 receptor complex. As with monocytes, signaling kinetic studies revealed that the observed differences in signaling output by the different IL-10 ligands were not a result of altered signaling activation kinetics (Figure 5e).

Granzyme B is a potent cytotoxic effector molecule that has been shown to be increased in CD8 T cells upon IL-10 stimulation (Naing et al., 2018). Next, we studied how granzyme B production by CD8 T cells was regulated by the different IL-10 ligands. For that, PBMCs or isolated CD8 T cells were activated following the workflow illustrated in Figure 6a and granzyme B levels were measured by flow cytometry or qPCR. As previously reported IL-10 stimulation did not affect classical early and late activation markers, i.e. CD69 and CD71 respectively, nor did it induce a significantly higher upregulation of inhibitory receptors, i.e. LAG-3 and PD-1 or affect CD8 cell proliferation (Sup. Figure 6a and 6b). On the other hand, IL-10 stimulation led to a strong upregulation of granzyme B levels in CD8 T cells both at the mRNA and protein levels, independently of whether CD8 T cells were activated in the context of a PBMC population or a purified CD8 T cell population (Sup. Figure 6c). When we compared our IL-10 ligands, at saturating concentrations WTD and R5A11D upregulated granzyme B production to a similar extent, 2.5-fold higher than granzyme B levels induced by TCR stimulation alone (Figure 5f). WTM showed very poor granzyme B production in agreement with its weak STAT activation. At a sub-saturating concentration, we again observed a stronger upregulation of granzyme B levels induced by R5A11D. R5A11M stimulation resulted in two major populations, with half of the donors upregulating granzyme B to levels similar to those induced by WTM and the other half upregulating granzyme B to levels comparable to those induced by the dimeric molecules. Overall, our results show that enhanced affinity for IL-10Rβ bestows IL-10 with robust activities over a wide range of ligand doses and immune cell subsets.

**Figure 6.**
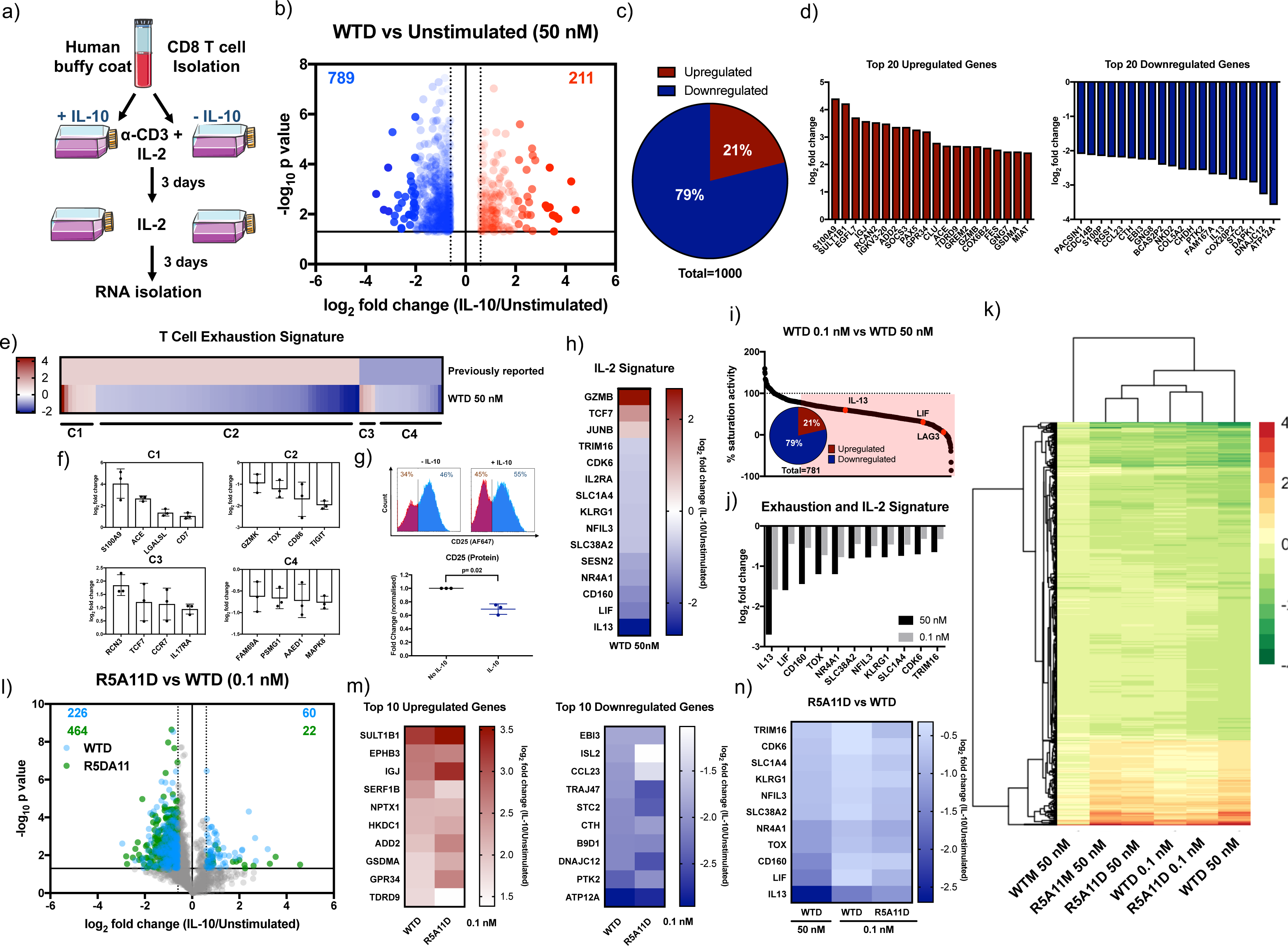
Characterisation of transcriptional activity induced by IL-10 and high affinity variants in human CD8 T cells. **(a).** Schematic of CD8 T cell stimulation. CD8 T cells were isolated by MACS and activated with anti-CD3/CD28 beads and IL-2 in the presence or absence of IL-10 wt and variants for three days. On day three the media was changed to IL-2 in the presence or absence of IL-10 wt and variants and cells were expanded for a further three days. **(b).** Volcano plot of CD8 T cell genes significantly upregulated by IL-10 wild type dimer ≧ 0.6 log_2_ fold change (red) and significantly downregulated ≦ −0.6 log_2_ fold change compared to non-IL-10 stimulated cells. Fold change was calculated by dividing WTD 50 nM by unstimulated values for each donor. The average fold change was calculated and the log_2_ of this value is plotted. P values were calculated by two-tailed unpaired t test of the log_2_ fold change of WTD 50 nM/unstimulated genes for each donor. Genes which were not significantly changed (>0.05) or were ≦ 0.6 ≧ −0.6 log_2_ fold change were excluded. **(c).** Proportion of CD8 T cell genes significantly regulated by WTD 50 nM ≧ 0.6 or ≦ −0.6 log_2_ fold change compared to unstimulated cells. **(d).** Log_2_ fold change for the top 20 protein coding genes significantly up (red) and down (blue) regulated by WTD 50 nM in CD8 T cells. **(e).** Heatmap of genes previously reported to be present in exhausted T cells. A list of exhaustion specific genes from (Bengsch et al., 2018) was used as a comparison for genes significantly up or down regulated by WTD 50 nM. Previously reported genes were given a nominal value of 1 for upregulated genes and −1 for downregulated genes. Log_2_ fold change for WTD 50 nM was plotted. Cluster 1 (C1) represents genes upregulated in exhausted cells and upregulated by WTD 50 nM. C2 represents genes upregulated in exhausted cells and downregulated by WTD 50nM. C3 represents genes downregulated in exhausted cells and upregulated by WTD 50 nM. C4 represents genes downregulated in exhausted cells and downregulated by WTD 50 nM. (f). The log2 fold change induced by WTD 50 nM for a sample of genes from each cluster is shown. Each point represents one biological replicate. **(g).** CD25 protein levels measured by flow cytometry. CD8 T cells were activated in a PBMC population for three days using anti-CD3 and IL-2 in the presence or absence of IL-10 (WTD, 50nM), followed by two days expansion with IL-2 in the presence or absence of IL-10. Fold change was calculated by dividing CD25 levels of IL-10 stimulated by non-IL-10 stimulated control values for each donor. Each point represents one biological replicate. P value was calculated by paired t test. **(h).** Heatmap showing the log_2_ fold change induced by WTD 50 nM for genes previously reported to be regulated by IL-2 in CD8 T cells (Rollings et al., 2018). **(i).** Percentage activity of low dose WTD compared to high dose WTD. The log_2_ fold change of WTD 0.1 nM was divided by WTD 50 nM and multiplied by 100. Genes which showed ≦ 75% of high dose activity (781 genes) are highlighted in red. Insert shows the percentage of these genes which up or downregulated activity. **(j).** Log_2_ fold change for genes associated with exhaustion or IL-2 stimulation and their regulation by WTD at 50 nM or 0.1 nM concentration. **(k).** Heatmap of genes significantly up or down regulated by WTD 50 nM ≧ 0.6 or ≦ −0.6 log_2_ fold change and their corresponding log_2_ fold change by R5A11D 50 nM, WTD 0.1 nM, R5A11D 0.1 nM, R5A11M 50 nM and WTM 50 nM compared to unstimulated control cells. Values are the mean of three donors. **(l).** Volcano plot of genes regulated by WTD (blue) and R5A11D (green) at 0.1 nM concentration each in CD8 T cells. Only genes which had already been showed to be significantly up or down regulated by WTD 50 nM are plotted. **(m).** Heatmap of the top 10 up and down regulated CD8 T cell genes by WTD 0.1 nM compared to R5A11D 0.1 nM. **(n).** Heatmap of exhaustion or IL-2 associated genes regulated by WTD at 50 nM and 0.1 nM and R5A11D 0.1 nM.

### Increased IL-10Rβ affinity enhances transcriptional activity of IL-10 in CD8 T cells

To obtain a more complete understanding of how IL-10 regulates CD8 T cells responses, we next performed transcriptional studies on CD8 T cells treated with the different IL-10 ligands. Human CD8 T cells were purified by positive selection and activated in the presence of WTD and our variants over 6 days as shown in Figure 6a. The transcriptional changes induced by WTD in CD8 T cells were less dramatic than those induced by this cytokine in monocytes. Only 1000 genes were significantly regulated, with 79% of those genes being down-regulated (Figure 6b and 6c). The most highly regulated genes are shown in Figure 6d. KEGG pathway analysis showed that IL-10 regulated genes are involved in cytokine-cytokine receptor interaction (Sup. Figure 7a). We noticed that IL-10 induced the downregulation of genes classically associated with CD8 T cell exhaustion (Figure 6e). We compared IL-10-regulated genes to a previously published list of exhaustion-specific CD8 T cell genes (Bengsch et al., 2018). Four clusters of exhaustion genes were identified that were regulated by IL-10. Cluster 1 comprises genes upregulated in both exhausted T cells and in T cells treated with IL-10. Cluster 2, the largest cluster, shows genes which were upregulated in exhausted T cells but downregulated by IL-10 treatment. Cluster 3 represent genes downregulated in exhausted T cells but upregulated by IL-10 treatment and cluster 4 is comprised of genes downregulated in both exhausted T cells and IL-10 treated T cells. A representative sample of regulated genes in each cluster is shown in Figure 6f. These results suggest that IL-10 may enhance CD8 T cell activities by preventing their exhaustion. We also observed a significant downregulation of IL-2R*α* by IL-10 treatment both at the mRNA (Sup. Figure 7b) and protein level (Figure 6g), which was associated with a reduction of expression of classical IL-2 dependent genes, such as IL-13, LIF, SLC1A4, NFIL3, etc (Figure 6h) (Rollings et al., 2018). Our results suggest that IL-10 regulates CD8 cytotoxic activities by limiting their sensitivity to IL-2, which may delay their exhaustion. As with monocytes, sub-saturating doses of WTD differentially affected a subset of genes regulated by IL-10, with the majority of those genes being downregulated by IL-10 treatment (Figure 6i). At sub-saturating doses WTD failed to regulate classical IL-2 dependent genes like IL-13 and LIF, suggesting that regulation of IL-2 activities by IL-10 requires high IL-10 doses (Figure 6j).

**Figure 7.**
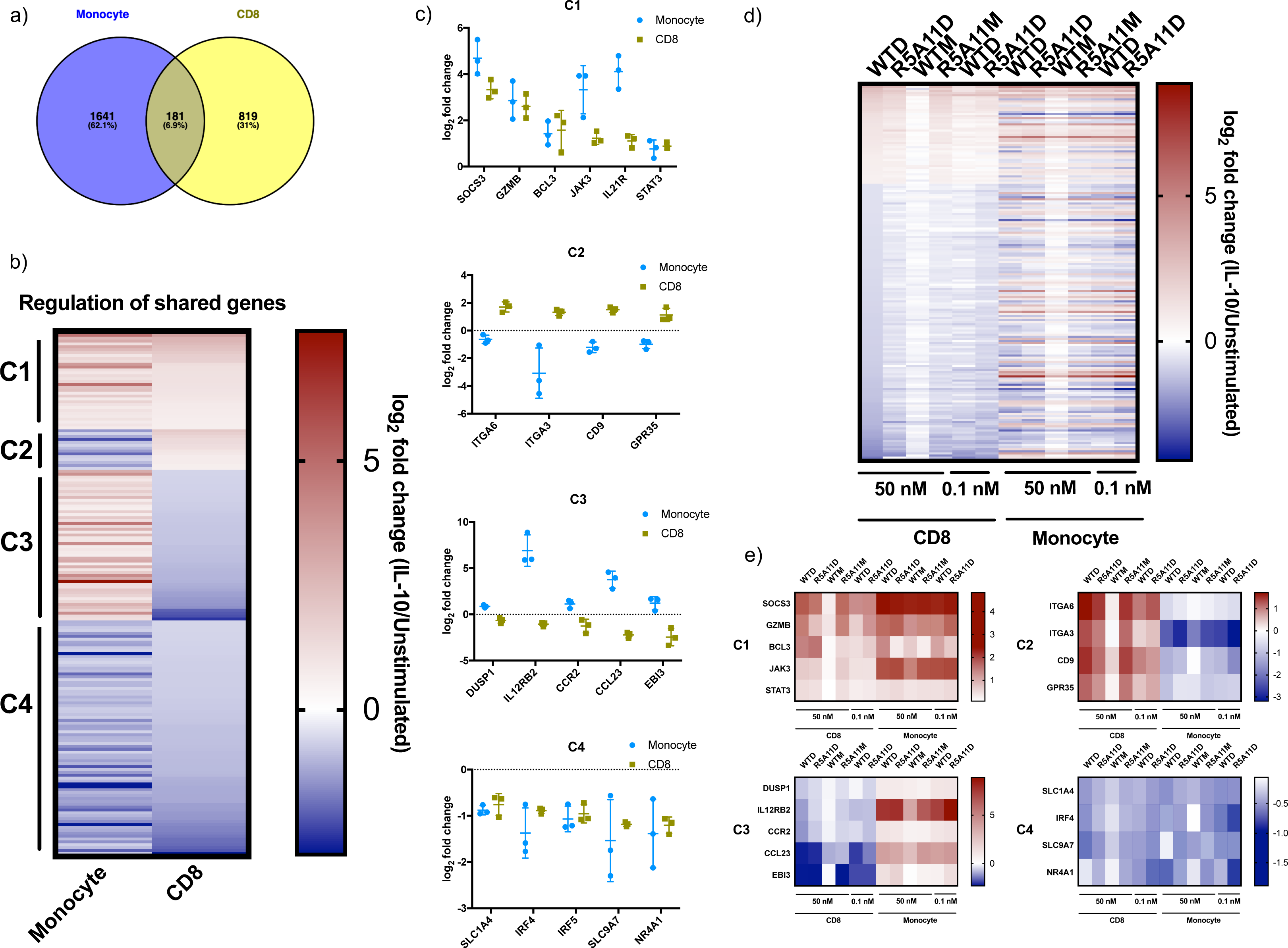
Comparison of common gene regulation by IL-10 in monocytes and CD8 T cells. **(a).** Venn diagram comparing genes significantly up or down regulated by wild type IL-10 (50 nM) in monocytes and CD8 T cells. Venn diagram generated using “Venny” (Oliveros, 2007-2015). **(b).** Comparison of the 181 genes regulated by IL-10 in both cell subsets. The log_2_ fold change for each gene of WTD (50 nM)/ unstimulated in both CD8 T cells and monocytes are plotted. Genes which are upregulated by IL-10 in both cell types are denoted as cluster 1 (C1). Genes upregulated by IL-10 in CD8 T cells but downregulated by IL-10 in monocytes are grouped in cluster 2 (C2). Genes which are upregulated by IL-10 in monocytes but downregulated by IL-10 in CD8 T cells are grouped in cluster 3 (C3). Genes downregulated by IL-10 in both monocytes and CD8 T cells are denoted by cluster 4 (C4). **(c).** Examples of genes from each cluster. The log_2_ fold change of WTD (50 nM)/ unstimulated in both CD8 T cells and monocytes are plotted. Each point represents one biological replicate and error bars represent the standard deviation. **(d).** Heatmap of the shared 181 genes and their expression in both cell subsets by IL-10 wild type and high affinity variants, monomers and dimers. The averge log_2_ fold change for each gene stimulated/unstimulated are plotted. **(e).** Regulation of genes from clusters 1-4 by the high affinity and monomeric variants.

As seen for monocytes, WTM showed very poor induction of gene expression (Figure 6k and Sup. Figure 7c), in line with its sub-optimal STAT activation. R5A11M again enhanced the transcriptional response when compared to WTM but failed to reach expression levels induced by the dimeric ligands despite activating very similar STAT signaling profiles (Figure 6k and Sup. Figure 7c). Indeed, when directly comparing expression levels induced by WTD and R5A11M, the high affinity monomer showed diminished activity of 56% of WTD regulated genes (Sup. Figure 7d). Similar to the results obtained with monocytes, R5A11D at 0.1 nM clustered with WTD 50 nM, supporting its ability to act effectively at low concentrations (Figure 6k). When the expression levels induced by WTD and R5A11D at the sub-saturating concentration were compared, we identified 38% of IL-10 high-dose regulated genes remained upregulated by R5A11D (Figure 6l and Sup. Figure 7e). This was clearly reflected when the expression of the top 10 up and downregulated genes by WTD and R5A11D at 0.1 nM were compared (Figure 6m). Importantly, classical IL-2 dependent genes, which were not regulated by WTD at low doses, were still regulated by R5A11D at these concentrations (Figure 6n). Together our data confirms that IL-10 variants with enhanced affinity for IL-10Rβ exhibit more robust activity at a wider range of ligand concentrations, which opens new avenues to boost IL-10 based anti-cancer immune-therapies.

### Differential gene expression program regulated by IL-10 in monocytes and CD8 T cells

Our study provides a highly detailed description of transcriptional changes induced by IL-10 in monocytes and CD8 T cells. Despite the obvious differencies in the manner that the two cell types were stimulated with IL-10, we investigated similarities of the transcriptional program induced by IL-10 in the two cell subsets, as a proxy to understand STAT3 transcriptional activities. To minimize variability resulting from the different treatments, we focused on genes that were regulated by IL-10 treatment in both monocytes and CD8 T cells. 181 genes were regulated by IL-10 in monocytes and CD8 T cells (Figure 7a). We could identify four gene clusters based on their regulation by IL-10 treatment (Figure 7b). Cluster 1 comprises genes that were upregulated by IL-10 treatment in both moncytes and CD8 T cells (Figure 7b). Cluster 2 corresponds to genes that were downregulated by IL-10 in monocytes, but upregulated by IL-10 in CD8 T cells. Cluster 3 show genes that were upregulated by IL-10 treatment in monocytes and downregulated by IL-10 treatment in CD8 T cells. Cluster 4 comprises genes downregulated by IL-10 treatment in monocytes and CD8 T cells. A representative sample of regulated genes in each cluster is shown in Figure 7c. The regulation of these shared genes was common to all IL-10 variants, with each variant modulating the induction (simply fold change relative to untreated) of these genes according to its receptor binding affinity (Figure 7d and 7e). Overall our comparative study highlights that IL-10 partially induces the regulation of a shared gene expression program between monocytes and CD8 T cells. However, whether those IL-10 regulated genes are induced or repressed by IL-10 treatment depends on the cellular context in which IL-10 stimulation takes place, providing an additional level of gene regulation by cytokines.

### R5A11D enhances the anti-tumor function of chimeric antigen receptor (CAR) T-cells

Chimeric antigen receptor (CAR) T cells have shown promising advances in cancer immune therapies in recent years. Thus, we determined whether the IL-10 variants would enhance the cytotoxic activities elicited by CAR T cells. For that, we used CAR T cell based model targeting acute myeloid leukemic cells (Warda et al., 2019). Human T cells were activated and transduced with anti-IL-1RAP-CD28--41BB-CD3ζ CAR or mock CAR (without anti-IL-1RAP) and initially expanded for 6 days in IL-2 conditioned media. After that initial expansion cells were cultured in IL-2 media contaning the WTD or R5A11D for an additional 3 days before performing cytotoxicity killing assay of target IL-1RAP expressing Mono-Mac-6 cells (Figure 8a and 8b). IL-1RAP CAR T cells cultivated in the presence of IL-2 +/− IL-10 variants showed specific cytotoxicity against Mono-Mac-6 cells in a ratio-dependent manner when compared to mock T cells (Figure 8c). Remarkably, at the different ratios tested, IL-1RAP CAR T cells cultured with IL-2/R5A11D exhibited higher levels of cytotoxicity than IL-1RAP CAR T cells culture with IL-2/WTD or IL-2 alone. While only 36% of Monomac-6 cells remained alive after incubation with IL-2/R5A11D treated IL-1RAP CAR T cells, 54% and 56% of Monomac-6 cells remained alive after incubation with IL-2/WTD or IL-2 IL-1RAP CAR T cells respectively. Collectively, our data demonstrate superior efficacy of R5A11D over IL-10 wt in boosting the cytotoxic activities of CAR T cells.

**Figure 8.**
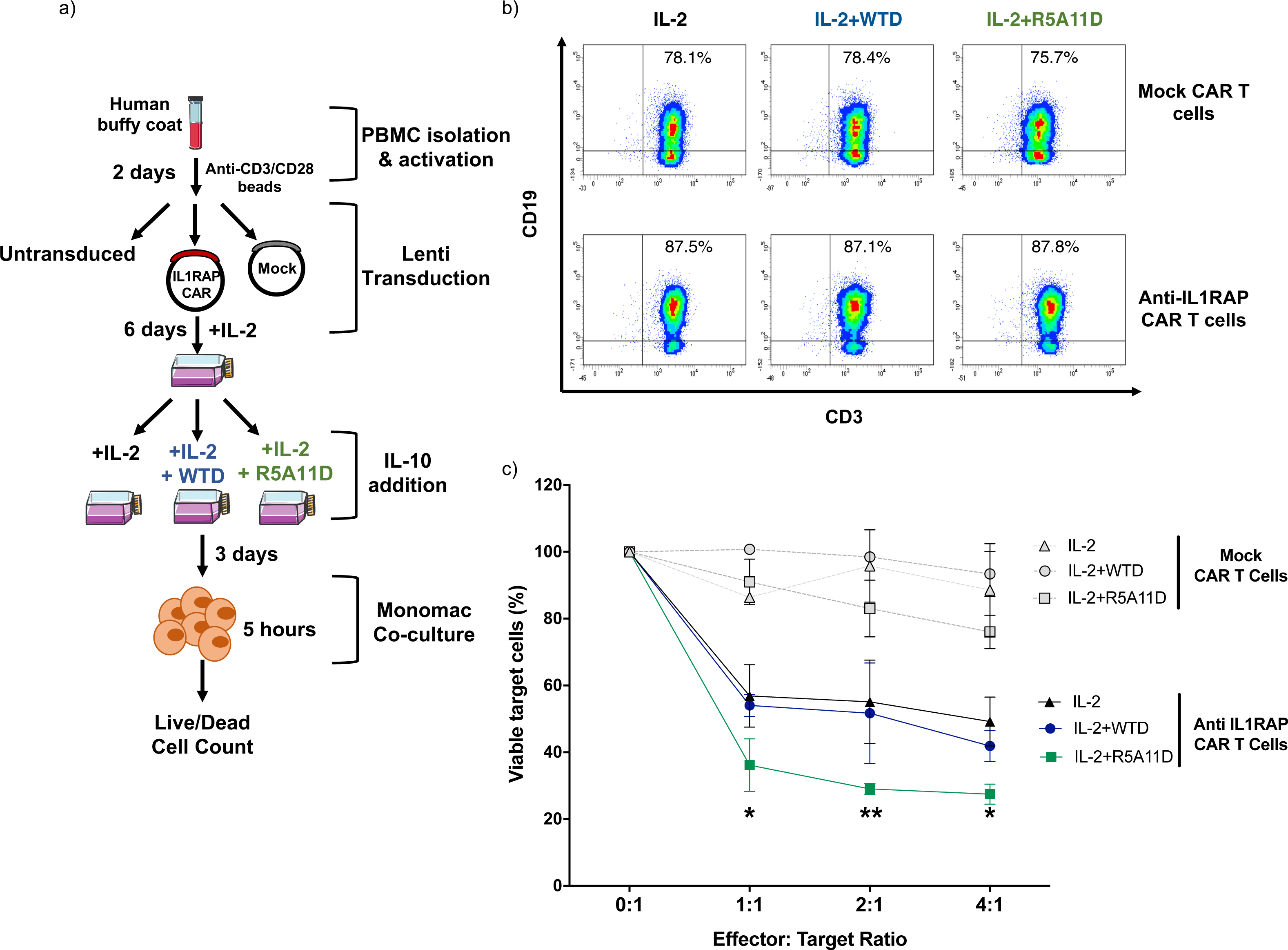
Evaluation of the effect of WTD and R5A11D to Chimeric Antigen Receptor T-cells (CAR T-cells) cytotoxicity. **(a)** Schematic of CAR T cells cytotoxic assay. Total T cells were isolated from PBMCs and activated with anti-CD3/CD28 dynabeads for 2 days. The T cells were then transduced with lentivirus supernatant. Effector T-cells (untransduced T-cells, Mock T-cells or IL-1RAP (**IL-1RAP:** Interleukin-1 Receptor Accessory Protein)-CAR T-cells) were cultivated in presence of IL-2 with or without WTD (25 nM) or R5A11D (25 nM) in the culture medium for 3 days. Then, the target cells, Mono-Mac-6 cells (IL-1RAP positive leukemic cell line), were co-cultured at mentioned Effector:Target (E:T) ratios. After 5 hours, the percentage of alive Mono-Mac-6 cells were measured by 7-AAD staining by flow cytometry. **(b)** Representative dot plots of transduced anti-IL-1RAP CAR expression profiles as analyzed by truncated CD19 marker expression on anti-IL-1RAP CAR T cells and Mock T cells cultured in the presence of IL-2 with or without WTD or R5A11D (25nM) for 3 days. **(c)** Target cells viability after 5 hours co-culture with effector cells. Data are presented as mean ±SEM (PBMCS from n=3 healthy donnors). *p<0.001, **p<0.0001 (IL-2 alone or IL-2+WTD vs. IL-2+R5A11D).

## DISCUSSION

IL-10 is an important immuno-modulatory cytokine that regulates inflammatory responses and enhances CD8 T cells cytotoxic activities (Moore et al., 2001; Oft, 2014; Walter, 2014). Despite its central role preserving immune homeostasis, there is still a dearth of knowledge of the exact molecular mechanisms through which IL-10 carries out its functions. We postulate that the weak binding affinity that IL-10 exhibits for IL-10R*β* critically contributes to its functional fitness, by limiting the range of concentrations at which IL-10 elicits its full immuno-modulatory potential. Here we have engineered IL-10 to enhance its affinity for IL-10R*β* to investigate whether the stability of the IL-10 receptor complex determines IL-10 bioactivity potencies. Two main findings arise from our study: (1) Affinity-enhanced IL-10 variants trigger more robust responses at a wide range of ligand concentrations and in different immune cell subsets than wildtype IL-10, and (2) the stoichiometry of the IL-10-receptor complex contributes to IL-10 bioactivity potencies beyond regulation of STAT activation levels. More generally, this work outlines a strategy to improve the potency of low receptor binding affinity cytokines and presents new molecular and cellular data with the potential to revitalise failed IL-10 therapies.

IL-10 exerted a profound regulation of the monocytic transcriptional program in our studies, agreeing with previous observations (Moore et al., 2001). IL-10 treatment inhibited antigen presentation by monocytes, limited their ability to recruit inflammatory immune cell subsets through regulation of chemokines and chemokine receptor expression, and boosted their phagocytic activity through the upregulation of scavenger receptors such as CD93, CD47, CD163 and cytokine receptors such as IL-21R*α*. In addition, IL-10 treatment modulated the metabolic activity of monocytes by altering their glycolytic and lipid biosynthesis potential, in line with recent studies (Ip et al., 2017). Interestingly, IL-10 effects were slightly biased towards gene repression, with 59% of genes regulated by IL-10 being downregulated. Indeed, several studies have reported the ability of STAT3 to inhibit transcription induced by other STATs (Costa-Pereira et al., 2002; Ray et al., 2014; Yang et al., 2011), suggesting that STAT3 activating cytokines may elicit their functions by disrupting transcriptional programs induced by other cytokines. In agreement with this model, we recently reported that IL-6, another STAT3 activating cytokine, promoted strong STAT3 binding to chromatin, but poor gene expression (Martinez-Fabregas et al., 2019).

The vast majority of reports in the literature describing IL-10 activities have focused on myeloid cells and use a single dose of IL-10, often at saturation (de Waal Malefyt et al., 1991a; Ding et al., 1993; Fiorentino et al., 1991a). However, we have a poor understanding regarding the range of IL-10 doses at which this cytokine elicits a full response in myeloid cells, a critical aspect when considering translation of this cytokine to the clinic. Here we provide transcriptional data from monocytes stimulated with two different doses of IL-10, one saturating and the second sub-saturating, with the latter more closely resembling the doses achieved during IL-10 therapies (Naing et al., 2018). Interestingly, 27% of genes regulated by IL-10 were affected when sub-saturating doses were used. The vast majority of affected genes (95%) were genes downregulated by IL-10 and encoded proteins critically contributing to establish an inflammatory environment i.e. key chemokines and cytokines such as IL-24, CXCL10, CXCL11, CCL22. This data suggests that IL-10 anti-inflammatory activities specifically require high and sustained doses to reach their full effect, explaining in part the failing of IL-10 therapies. Our engineered IL-10 variant exhibited a more robust activity at sub-saturating doses and induced potent inhibition of pro-inflammatory chemokines and cytokines, i.e. IL-24, CXCL10, CXCL11, CCL22. It is thus tempting to speculate that our engineered variant could rescue failed IL-10 therapies by promoting anti-inflammatory activities at low ligand doses.

The anti-inflammatory activities elicited by IL-10 and its effects on monocytes and macrophages are very well documented. How IL-10 regulates the activity of CD8 T cells on the other hand is less clear and more controversial (Oft, 2014). While some studies have reported that IL-10 enhances the function of CD8 T cells and their ability to kill tumour cells (Emmerich et al., 2012), others report that the presence of IL-10 in the tumour microenvironment predicts poor responses by inhibiting T cell activation (Zhao et al., 2015). Our results agree with a positive effect of IL-10 treatment on CD8 T cells cytotoxic activities. CD8 T cells stimulated in the presence of IL-10 exhibited enhanced levels of effector molecules such as granzyme B, agreeing with recent clinical trials that show an improvement in the tumour response of patients treated with Pegylated-IL-10 (Naing et al., 2019). In addition, our engineered IL-10 variants improved the killing activity of CAR-T cells *in vitro*. However, the molecular bases by which IL-10 boosts the anti-tumour CD8 T cell response remains poorly defined. Our transcriptional studies highlighted that CD8 T cells stimulated with IL-10 exhibited a reduced exhaustion gene signature and were more functionally fit. IL-10 treated CD8 T cells also expressed lower levels of IL-2R*α*, which correlated with a reduced IL-2 gene signature in these cells. Alltogether, our data agree with a model where IL-10 may reduce the sensitivity of CD8 T cells to IL-2 and their transition towards an exhausted phenotype. Our engineered IL-10 variant outperformed IL-10 wildtype in every read out tested when sub-saturating doses were used, reproducing our observations in monocytes and highlighting its potential to boost anti-tumour responses at therapeutical doses.

The importance of the dimeric IL-10 architecture for generating its biological responses is not yet well understood. WTD binds IL-10R*α* 60-fold stronger avidly than WTM, which contributes to its more efficient recruitment of IL-10R*β* to the signaling complex and its more potent activities (Tan et al., 1993). Paradoxically, R5A11M, which binds IL-10R*β* with higher affinity and elicits receptor assembly levels similar to WTD, triggers weaker transcriptional responses, despite activating STATs to a very similar extent as WTD. In addition, viral IL-10 (also a dimeric ligand) induces the same specific activity as WTD even though it binds IL-10R*α* with lower affinity than WTM (Liu et al., 1997; Yoon et al., 2005). Overall these observations suggest that in addition to receptor binding affinity, the stoichiometry of the IL-10-receptor complex contributes to fine-tune IL-10 bioactivity potencies. We recently showed that the number of phospho-tyrosines available in cytokine receptor intracellular domains critically contribute to defining signaling identity by cytokines (Martinez-Fabregas et al., 2019). IL-6 variants that triggered partial phosphorylation of Tyr available in the gp130 intracellular domain exhibited a biased STAT3 versus STAT1 activation (Martinez-Fabregas et al., 2019). A similar model could be invoked to explain functional differences between monomeric and dimeric IL-10 ligands. Different patterns of phospho-Tyr in IL-10R*α* and IL-10R*β* could be induced in the context of the hexameric receptor complex formed by the dimeric ligands, which could trigger the activation of additional signaling pathways and provide functional specificity. In agreement with this model, WTM and R5A11M elicited biased STAT3 activation in CD8 T cells. Future studies will be needed to address whether the higher number of Tyr available in the hexameric complex engaged by WTD contribute to define its signaling signature and biological identity.

## ACKNOWLEDGMENTS

We thank members of the Moraga, Mitra, Walter and Piehler laboratories for helpful advice and discussion. This work was supported by the Wellcome Trust 203752/Z/16/Z (C.G.), by the Wellcome-Trust-202323/Z/16/Z and ERC-206-STG grant (I.M.), by EMBO (S.W. 454-2017) and by the DFG (SFB 944, P8/Z, J.P.) and NIH (R01 AI143554, M.R.W.).

## COMPETING INTERESTS

The authors declare that they have no competing interests.

## MATHERIALS AND METHODS

### Protein expression and purification

Monomeric wild type IL-10 (Josephson et al., 2000), monomeric high affinity variants and IL-10Rα ectodomain (amino acids 22-235) were cloned and expressed as described in (Martinez-Fabregas et al., 2019). Briefly, protein sequences were cloned into the pAcGP67-A vector (BD Biosciences) in frame with an N-terminal gp67 signal sequence, driving protein secretion, and a C-terminal hexahistidine tag. The baculovirus expression system was used for protein production as outlined in (LaPorte et al., 2008). *Spodoptera frugiperda* (SF9) cells, grown in SF900II media (Invitrogen), were transfected to produce P_0_ baculovirus stocks that were then expanded in SF9 cells to produce P_1_ virus stock. Protein expression was performed using *Trichoplusiani ni* (High Five) with cells grown in InsectXpress media (Lonza).

Purification was performed using the method described in (Spangler et al., 2019). Briefly, the cells were pelleted with centrifugation at 2000 rpm, prior to a precipitation step through addition of Tris pH 8.0, CaCl_2_ and NiCl_2_ to final concentrations of 200mM, 50mM and 1mM respectively. The precipitate formed was then removed through centrifugation at 6000 rpm. Nickel-NTA agarose beads (Qiagen) were added and the target proteins purified through batch binding followed by column elution in HBS, 200mM imidazole, pH 7.2. Target proteins were concentrated and further purified by size exclusion chromatography on an ENrich SEC 650 300 column (Biorad), equilibrated in 10 mM HEPES (pH 7.2), 150 mM NaCl. IL-10Rα was biotinylated using EZ-Link NHS biotinylation kit (Thermo) according to the manufacturer’s protocols.

For expression of biotinylated IL-10Rβ the ectodomain (amino acids 20-220) was cloned into the pAcGP67-A vector carrying a C-terminal biotin acceptor peptide (BAP)-LNDIFEAQKIEWHW followed by a hexahistidine tag. The purified protein was biotinylated with BirA ligase following the protocol described in (Spangler et al., 2019).

For expression of dimeric wild type IL-10 and dimeric high affinity variants, synthesised gene blocks (IDT) were cloned into the pET21 vector in frame with an N-terminal hexahistidine tag and a lac promotor, and transformed into *E. Coli* BL21 cells. Protein production was induced using 1 mM final concentration of IPTG (Formedium) followed by incubation at 37°C for 3 to 5 hours. Cells were harvested by centrifugation at 6000 xg for 15 minutes. The cell pellets were resuspended in 50 mM Tris-HCl (pH 8.0), 25% (w/v) sucrose, 1 mM Na EDTA, 10 mM DTT, 0.2 mM PMSF per litre of original culture and frozen at −80°C overnight.

The recombinant protein was expressed as inclusion bodies, purification of which was performed as follows. Cell pellets were lysed in 100 mM Tris-HCl (pH 8.0), 2% (v/v) TritonX-100, 200 mM NaCl, 2500 units Benzonase, 10 mM DTT, 5 mM MgCl2, 0.2 mM PMSF and incubated for 20 minutes with stirring at room temperature. 10 mM EDTA final concentration was then added to the suspension and the cells were sonicated (8-10 cycles of 15 seconds on/off, 15 microns, Soniprep 150) in an ice bath. The solution was centrifuged at 7000 xg for 15 mins (4°C) and the pellet was resuspended in 50mM Tris-HCl pH 8.0, 0.5% Triton X-100, 100 mM NaCl, 1 mM Na EDTA, 1 mM DTT, 0.2 mM PMSF. This step was repeated for a total of at least three washes until the preparation appeared white. The final pellet was then washed once in detergent free buffer (50 mM Tris-HCl pH 8.0, 1 mM Na EDTA, 1 mM DTT, 0.2 mM PMSF).

The purified inclusion bodies were solubilised in 10 mls of 6M GuHCl per litre of original culture, for 30 minutes at room temperature. The solution was clarified by a centrifugation at 7000 rcf for 15 minutes and the solubilised protein was carefully decanted. Refolding was performed through dropwise addition of the solubilised protein solution into refolding buffer (50 mM Tris-HCl, pH 8.0, 50 mM NaCl, 5 mM EDTA, 2 mM reduced glutathione (GSH) and 0.2 mM oxidized glutathione (GSSG)) at a ratio of 1:20 solution:buffer at 4°C followed by incubation with gentle stirring overnight at 4°C.

The solution was then filtered to remove any precipitant and dialysis performed against 10 mM HEPES (pH 7.2),150 mM NaCl, using dialysis membrane with a 14 kDa Mwt cut off.

After dialysis protein was then further purified using Ni-NTA beads and by size exclusion on a Superdex75 increase 10/300 column (GE Healthcare). Endotoxin removal was then performed as follows. 1 mL of Ni-NTA agarose was added to a polyprep column and equilibrated with 10 mls of HBS before addition of the protein. The column was washed with 50 column volumes of ice-cold HBS, 150 mM NaCl, 20 mM imidazole, 0.1% Triton-X114 (pH 7.2) to remove endotoxin. The column was then washed with a further 20 column volumes of HBS, 20 mM imidazole (pH 7.2). The now endotoxin-free protein was eluted using 4 column volumes of HBS, 200 mM imidiazole (pH 7.2). The protein was buffer exchanged into 10 mM HEPES, 150 mM NaCl (pH 7.2), using PD-10 columns (GE Healthcare). Endotoxin levels were measured using Pierce LAL Chromogenic Endotoxin Quantitation Kit (Thermo) following the manufacturer’s protocol. For all proteins endotoxin levels were below detection levels of the kit.

### Surface Plasmon Resonance

Surface plasmon resonance was used to determine the binding affinity of the recombinantly produced monomeric IL-10 wild type and variants to IL-10Rβ in the presence or absence of IL-10Rα. Biotinylated IL-10Rβ was immobilised onto the chip surface via streptavidin. Series S Sensor SA (GE Healthcare) chips were primed in 10 mM HEPES, 150 mM NaCl, 0.02% TWEEN-20, prior to immobilisation of the biotinylated receptor. Analysis runs were then performed in 10 mM HEPES, 150 mM NaCl, 0.05% TWEEN-20 and 0.5% BSA. A Biacore T100 (T200 Sensitivity Enhanced) was used for measurement with Biacore T200 Evaluation Software 3.0 used for data analysis.

### Cell culture

Human buffy coats were obtained from the Scottish Blood Transfusion Service and peripheral blood mononuclear cells (PBMCs) were isolated by density gradient centrifugation (Lymphoprep, StemCell Technologies). PBMCs were grown in RPMI-1640, 10% v/v FBS, 100 U/mL penicillin-streptomycin (Gibco) and cytokines for proliferation and activation as follows. For three days media was supplemented with 100 ng/mL anti-CD3 (human UltraLEAF, Biolegend) and 20 ng/mL IL-2 (Proleukin, Novartis) in the absence or presence of IL-10 variants. After three days of activation cells were centrifuged and resuspended in media supplemented with 20 ng/mL of IL-2 plus or minus IL-10 variants. Cell populations were allowed to expand for 2-3 days.

Monocytes were isolated from PBMC populations using CD14 positive selection. Anti-CD14^FITC^ antibody (Biolegend #367116) was used to stain cells and isolation was done by magnetic separation following manufacturer’s protocol (MACS Miltenyi). Monocytes were then cultured in complete RPMI (as above) supplemented with M-CSF (20 ng/mL, Biolegend). Cells were then stimulated with IL-10 variants for twenty-four hours before analysis.

CD8 T cells were isolated from PBMCs by magnetic separation (MACS Miltenyi) after staining with anti-CD8a^FITC^ antibody (Biolegend #30906). For activation of purified CD8 T cells ImmunoCult Human CD3/CD28 T cell Activator (Stem Cell) was used following manufacturer’s protocol as well as the addition of 20 ng/mL IL-2 and IL-10 variants. Cells were activated for 3 days and then the media was replaced with complete RPMI supplemented with 20 ng/mL IL-2 as well as IL-10 variants for 2-3 days.

### CAR-T cytotoxicity assay

Lentiviral construction and genetically modified Chimeric Antigen Receptor (CAR) T-cell transduction: The Interleukin-1 Receptor Accessory Protein (IL-1RAP) (Interleukin-1 Receptor Accessory Protein) CAR lentiviral construct was generated as previously described(Warda et al., 2019)1. Briefly, this vector carries a 3rd generation CAR, an inducible Caspase 9 (iCASP9) (inducible Caspase 9) suicide gene and a delta CD19 surface gene. The Mock control vector doesn’t contain the CAR sequence.

Peripheral blood mononuclear cells (PBMCs) were isolated by Ficoll gradient density centrifugation using Ficoll-Paque (Velizy-Villacoublay, France) with anonymous blood samples collected from healthy donors at a French blood center (Besançon, France). Donors provided written informed consent; the study was conducted in accordance with the ethical guideline (declaration of Helsinki) and approved by the local ethical the CPP-Est committee (France). Human tumor Mono-Mac-6 cell line was obtained from DSMZ German collection of microorganisms and cell culture GmbH and stored in a master cell bank. The cells were cultivated in RPMI-1640 supplemented with 10% FBS, penicillin-streptomycin.

CD3/CD28 activated cells were established from healthy donors and transduced with lentiviral supernatant encoding the IL-1RAP CAR or Mock sequences. At day 6 post-transduction, cells were put in the different culture conditions: IL-2 (500 UI/mL) with or without WTD or R5A11D at 25nM for 3 days. IL-1RAP CAR T-cells, Mock T-cells and untransduced T-cells were then used for the functionally cytotoxic tests.

For CAR T-cell cytotoxicity assay, prior to the co-culture, effector cells (untransduced T-cells, Mock T-cells and IL-1RAP CAR T-cells) were labeled with e-FluorTM v450 (eBioscience, #65-0842-85) in order to differenciate them from tumor cells. Then effector cells were co-cultured with Mono-mac-6 cells at different Effector:Target (E:T) ratios (1:1; 2:1 and 4:1). After 5 hours, cells were stained with 7-Amino-Actinomycin D (7-AAD; BD Bioscience, #559925) and the percentage of alive Mono-Mac-6 cells was determined by flow cytometry using a FACSCanto II flow cytometer (BD Bioscience).

### Flow cytometry staining and antibodies

For live cell surface staining of HLA-DR^PE^ (Biolegend #307605) non-adherent monocytes were removed from culture by centrifugation and resuspension in cold PBS. Adherent monocytes were detached using Acutase (StemCell Technologies) at room temperature for 5 to 10 minutes. Cells were kept at 4°C or on ice during live cell surface marker staining and staining was done in 96-well v-bottom plates (Griener) unless otherwise stated. Non-adherent and detached cells were combined and resuspended in FcR blocking reagent (Miltenyi) for 10 minutes at 4°C in a volume of 50 µL per condition. Cells were washed in PBS/0.5% BSA and resuspend in 50 µL of antibody mixture diluted 1/100 in FcR blocking reagent. Antibody incubation was done for 30 to 60 minutes at 4°C in the dark. Cells were washed twice before resuspension in 100 µL per well for analysis on the CytoFlex flow cytometer (Beckman Coulter). Mean fluorescence intensity (MFI) was quantified for all populations. Data was normalised within each donor by dividing MFI of IL-10 treated cells by a non-IL-10 treated control from the same donor to calculate fold change.

For granzyme B intracellular staining either PBMCs or CD8 cells on day 6 of activation were fixed with 2% paraformaldehyde for 10 minutes at room temperature before washing in PBS. Cells were permeabilised in 0.1% Triton-X100/PBS for 10 minutes and washed in PBS/0.5% BSA. Cells were stained with anti-CD8a^AlexaFluor700^ (Biolegend #300920), anti-CD4^PE^ (Biolegend #357404), anti-CD3^BrilliantViolet510^ (Biolegend #300448) and anti-granzyme B^FITC^ (Biolegend #515403) at 1/100 dilution in PBS/0.5% BSA for one hour before washing. MFI was quantified for all populations and normalisation was done as described above.

For phospho-flow analysis of STAT1 and STAT3 cells were plated at 50 μL of cell suspension per well at a density of 2 × 10^5^ cells per well in 96-well V bottom plates. For dose response studies cells were simulated with 7-fold serially diluted IL-10 variants and an unstimulated control (50 μL per well) for 15 minutes at 37°C before fixation with 2% paraformaldehyde for 10 minutes at room temperature. For kinetic studies cells were stimulated with a saturating concentration of IL-10 variants (50 nM) at defined time points before fixation simultaneously with 2% paraformaldehyde. Cells were washed in PBS and permeabilised in ice-cold 100% methanol and incubated on ice for a minimum of 30 minutes. Cells were fluorescently barcoded as described in (Krutzik and Nolan, 2006; Martinez-Fabregas et al., 2019). Briefly, a panel of 16 combinations of two NHS-dyes (Pacific Blue and DyLight800, Thermo) were used to stain individual wells on ice for 35 minutes before stopping the reaction by washing in PBS/0.5% BSA. Once barcoded the 16 populations were pooled together for antibody staining. PBMCs, CD8 cells and monocytes were stained with the cell surface markers described above as well as anti-pSTAT3^Alexa488^ (Biolegend #651006) and anti-pSTAT1^Alexa647^ (Cell Signaling Technologies #8009). During acquisition individual populations were identified according to the barcoding pattern and pSTAT3^Alexa488^ and pSTAT1^Alexa647^ MFI was quantified for all populations. MFI was plotted and sigmoidal dose response curves were fitted using Prism software (Version 7, GraphPad). Data was normalised by assigning the highest MFI of the top concentration of all stimuli as 100% and the lowest MFI as 0% within each donor group.

### Yeast display library

Yeast surface display protocol was adapted from previous protocols (Boder and Wittrup, 1997; Martinez-Fabregas et al., 2019). To create an IL-10 yeast display library the monomeric IL-10 gene (Josephson et al., 2000) was subject to error-prone PCR as described in (Mendoza et al., 2017). This product was then amplified and transformed along with a linearized pCT302 vector into the *Saccharomyces cerevisiae* strain EBY100 and grown in selective dextrose casamino acids (SDCAA) media at 30°C for two days. Yeast cells were then place in selective galactose casamino acids (SGCAA) at 20°C for two days to induce cell surface expression of IL-10 variants as described in (Chao et al., 2006). Magnetic activated cell sorting (MACS, Miltenyi) was used to select for IL-10 variants with increased binding affinity for IL-10Rβ as described previously for other systems (Moraga et al., 2015b). Briefly, the first round of selection was performed using high concentrations of streptavidin beads to remove any yeast which displayed variants capable of binding streptavain. The second round of selection selected for yeast which display variants with the c-myc tag at their C-terminus, ensuring that displayed proteins were properly folded. The subsequent rounds of selection were carried out by incubating induced yeast with decreasing concentrations of recombinantly produced biotinylated IL-10Rβ for 2 hours followed by a 15 minute incubation with fluorescently labelled streptavidin (AlexaFluor647). Magnetic activated cell sorting (MACS, Miltenyi) selected for yeast which displayed IL-10 variants capable of binding IL-10Rβ. Once the concentration of IL-10Rβ needed for binding was decreased sufficiently compared to wild type monomeric IL-10, the yeast were plated on SDCAA agar and single colonies were isolated for dose response studies to determine the EC50 values of the mutants.

Yeast colonies displaying promising IL-10 variants were subject to Zymoprep (ZymoResearch) to isolate the plasmid which was then heat shocked into competent DH5α *E. coli*. These were subject to miniprep (Promega) and plasmids were sequenced to observe where mutations had occurred in the monomeric IL-10 gene. These genes were then cloned into the baculovirus expression vector pACgp67BN and recombinantly expressed as described above.

### Measurement of IL-6 secretion

Monocytes were stimulated with LPS (100 ng/mL) (*E. coli* O26:B6, Sigma) plus IL-10 variants at various concentration for 8 hours. Supernatant was then removed and used for enzyme linked immunosorbent assay (ELISA) for IL-6 detection (Biolegend, #430501). Manufacturer’s protocol was followed. 96-well half-area plates (Sigma) were coated in capture antibody and incubated overnight at 4°C. Plates were washed in PBS/0.05% Tween-20 and blocked for 1 hour in assay diluent and washed. Supernatant was diluted 1 to 10 in assay buffer before addition to the plate. The plates were incubated at room temperature for two hours with shaking. Plates were washed again and incubated for 1 hour with detection antibody. After washing, avidin-HRP was added and incubated for 30 minutes followed by incubation with TMB substrate solution for 15 minutes. The reaction was stopped by addition of H_2_SO_4_ and absorbance was measured at 450 nm and 570 nm with absorbance at 570 nm being subtracted from 450 nm.

### RNA Transcriptome Sequencing

Human primary monocytes and CD8 T cells from three donors each (StemCell Technologies) were stimulated as described in above. Cells were washed in Hank’s balanced salt solution (HBSS, Gibco) and snap frozen for storage. RNA was isolated using the RNeasy Kit (Quiagen) according to manufacturer’s protocol. All RNA 260/280 ratios were above 1.9. 1 μg of RNA was used per sample. Transcriptomic analysis was done by Novogene as follows. Sequencing libraries were generated using NEBNext® UltraTM RNALibrary Prep Kit for Illumina® (NEB, USA) following manufacturer’s recommendations and index codes were added to attribute sequences to each sample. Briefly, mRNA was purified from total RNA using poly-T oligo-attached magnetic beads. Fragmentation was carried out using divalent cations under elevated temperature in NEBNext First StrandSynthesis Reaction Buffer (5X). First strand cDNA was synthesized using random hexamer primer and M-MuLV Reverse Transcriptase (RNase H-). Second strand cDNA synthesis was subsequently performed using DNA Polymerase I and RNase H. Remaining overhangs were converted into blunt ends via exonuclease/polymerase activities. After adenylation of 3’ ends of DNA fragments, NEBNext Adaptor with hairpin loop structure were ligated to prepare for hybridization. In order to select cDNA fragments of preferentially 150∼200 bp inlength, the library fragments were purified with AMPure XP system (Beckman Coulter, Beverly, USA). Then 3 μl USER Enzyme (NEB, USA) was used with size-selected, adaptor-ligated cDNA at 37 °C for 15 min followed by 5 min at 95 °C before PCR. Then PCR was performed with Phusion High-Fidelity DNA polymerase, Universal PCR primers and Index (X) Primer. At last, PCR products were purified (AMPure XP system) and library quality was assessed on the Agilent Bioanalyzer 2100 system.

### RNA Sequencing Data Analysis

Primary data analysis for quality control, mapping to reference genome and quantification was conducted by Novogene as outlined below.

Quality control: Raw data (raw reads) of FASTQ format were firstly processed through in-house scripts. In this step, clean data (clean reads) were obtained by removing reads containing adapter and poly-N sequences and reads with low quality from raw data. At the same time, Q20, Q30 and GC content of the clean data were calculated. All the downstream analyses were based on the clean data with high quality.

Mapping to reference genome: Reference genome and gene model annotation files were downloaded from genome website browser (NCBI/UCSC/Ensembl) directly. Paired-end clean reads were mapped to the reference genome using HISAT2 software. HISAT2 uses a large set of small GFM indexes that collectively cover the whole genome. These small indexes (called local indexes), combined with several alignment strategies, enable rapid and accurate alignment of sequencing reads.

Quantification: HTSeq was used to count the read numbers mapped of each gene, including known and novel genes. And then RPKM of each gene was calculated based on the length of the gene and reads count mapped to this gene. RPKM, (Reads Per Kilobase of exon model per Million mapped reads), considers the effect of sequencing depth and gene length for the reads count at the same time and is currently the most commonly used method for estimating gene expression levels.

Statistical analysis was done by the authors in Excel. The fold change was calculated by dividing the IL-10 stimulated expression levels by the unstimulated control within each donor. The average fold change was calculated for each stimulation across the three donors and the log_2_ of this average was then calculated. For calculation of significantly changed genes, the log_2_ of the fold change between IL-10 stimulated and unstimulated expression levels of each donor was calculated, separately and an unpaired, two tailed t test was used to generate the p value. The log_10_ of this p value was then plotted against the previously calculated log_2_ average fold change. Genes which were significantly (p≦ 0.05) changed greater than 0.6 or less than −0.6 log_2_ fold change in the wild type IL-10 dimer (WTD) 50 nM condition were taken as a set list of genes against which all other IL-10 stimulations were compared. Upregulated genes were denoted as genes ≧ 0.6 log_2_ fold change and downregulated genes were denoted as genes ≦ −0.6 log_2_ fold change. For comparison of WTD to other IL-10 variant stimulations the average log_2_ fold changes of the variants were divided by the average log_2_ fold change of WTD. Genes with an RPKM of less than 1 in two or more donors were excluded from analysis so as to remove genes with abundance near detection limit.

Functional annotation of genes (KEGG pathways, GO terms) was done using DAVID Bioinformatics Resource functional annotation tool (Huang da et al., 2009a, b). Clustered heatmap was generated using R Studio Pheatmap package.

### Live-cell dual colour single molecule imaging studies

Receptor homo- and heterodimerization was quantified by two-colour single-molecule co-tracking as described previously (Moraga et al., 2015c; Wilmes et al., 2015; Wilmes et al., 2020). Receptor dimerization experiments were performed in HeLa cells transienty expressing IL-10Rα and IL-10Rβ with N-terminally fused variants of monomeric ECFP and EGFP, respectively. Cell surface Labelling was achieved using anti-GFP nanobodies Minimizer(MI) and Enhancer(EN), respectively, site-specifically conjugated with photostable fluorophores via an engineered cysteine residue. For quantification of receptor heterodimerization, IL-10Rα and IL-10Rβ were labelled with MI^Rho11^ (ATTO Rho11, ATTO-TEC GmbH) and EN^AT643^ (ATTO 643, ATTO-TEC GmbH), respectively. For quantification of homodimerization, either IL-10Rα was labelled with MI^Rho11^ and MI^AT643^, or IL-10Rβ was labelled with EN^Rho11^ and EN^AT643^. Over-expression of the corresponding other receptor subunit was ensured by labelling with EN^AT488^ or MI^AT488^ (ATTO 488, ATTO-TEC GmbH), respectively. Time-lapse dual-color imaging of individual IL-10Rα and IL-10Rβ in the plasma membrane was carried out by total internal reflection fluorescence microscopy with excitation at 561 nm and 640 nm and detection with a single EMCCD camera (Andor iXon Ultra 897, Andor) using an image splitter (QuadView QV2, Photometrics). Molecules were localized using the multiple-target tracing (MTT) algorithm (Serge et al., 2008). Receptor dimers were identified as molecules that co-localized within a distance threshold of 150 nm for at least 10 consecutive frames as described in detail previously (Moraga et al., 2015c; Wilmes et al., 2015; Wilmes et al., 2020).

---

**Supplementary Figure 1.**
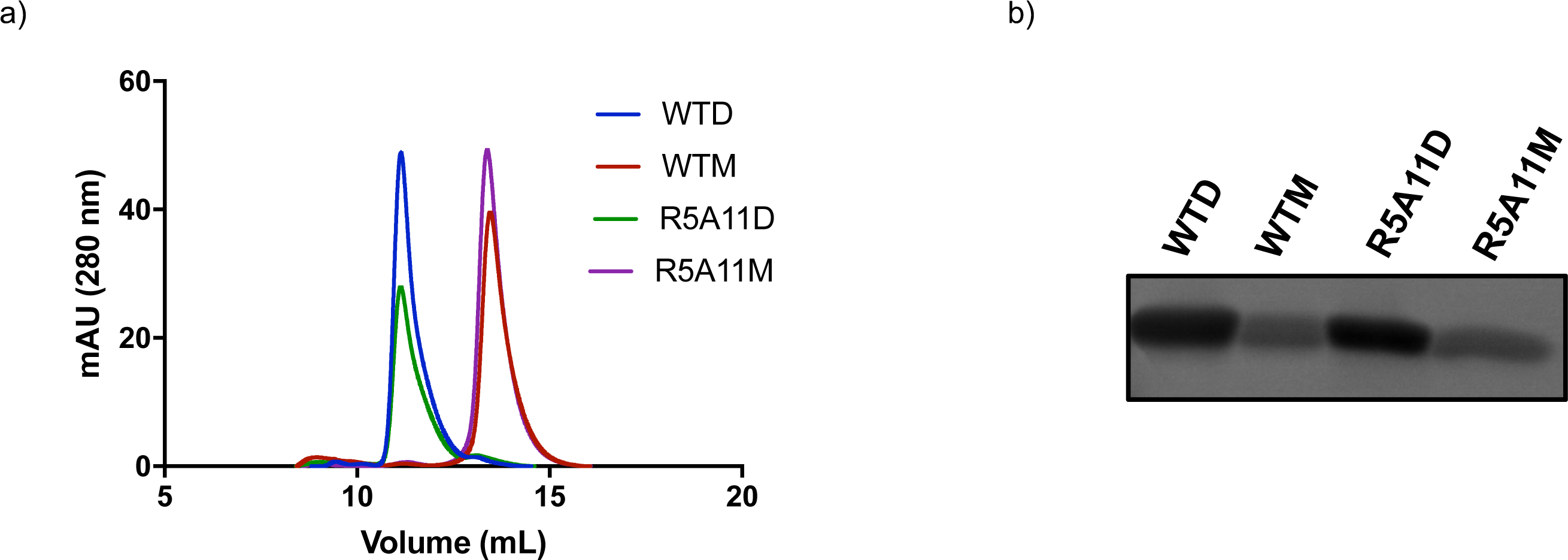
Recombinant expression of wild type and high affinity IL-10 monomeric and dimeric variants. **(a).** FPLC chromatogram for wild type and mutant monomer and dimer. Proteins were run on an S200 gel filtration column and separation by size exclusion. **(b).** Coomassie gel of FPLC purified proteins run on 10% gel.

**Supplementary Figure 2.**
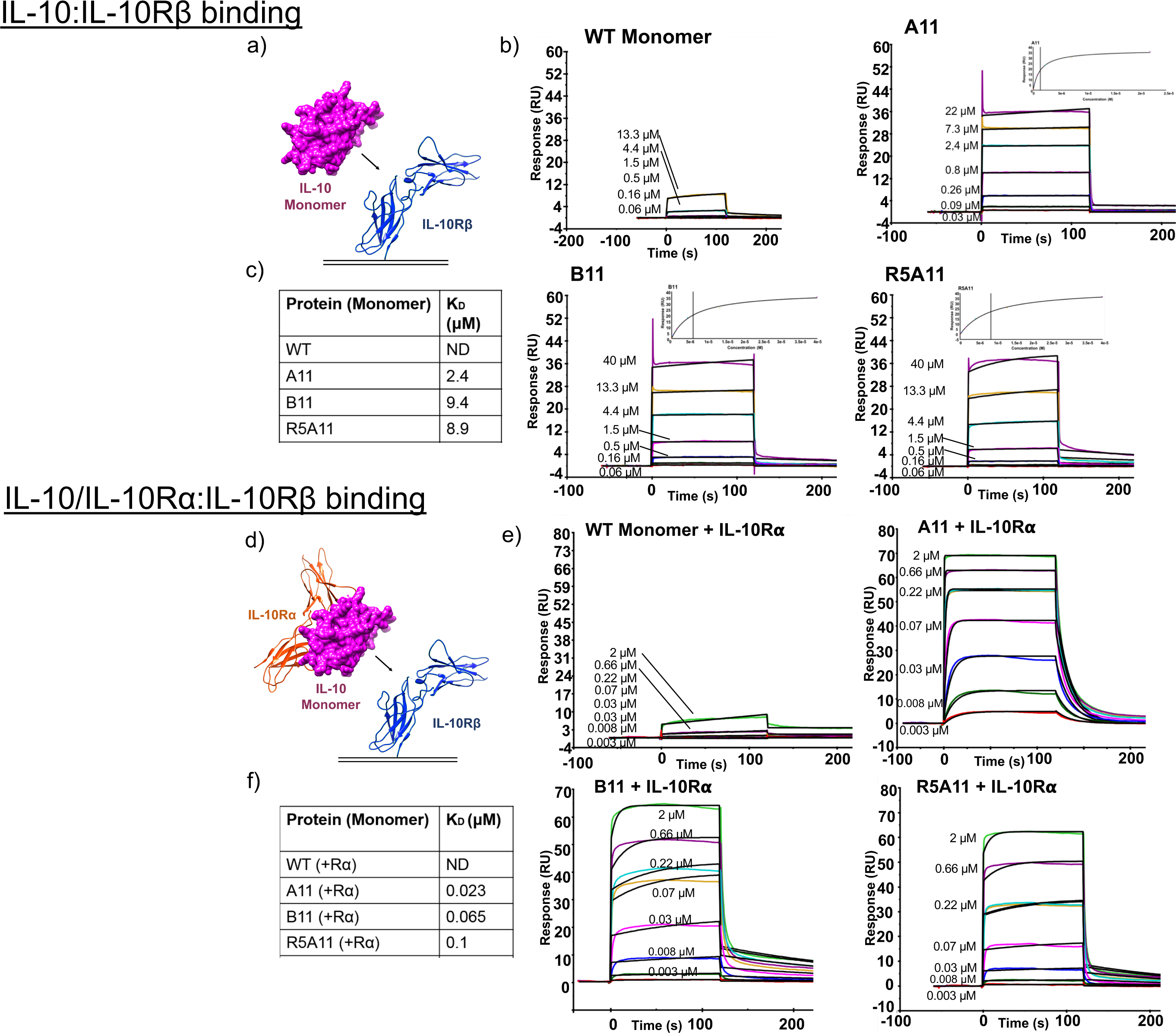
Biophysical characterisation of high affinity IL-10 variants. **(a).** For biacore measurement IL-10Rβ is immobilised on the chip surface via biotin-streptavidin interaction and IL-10 variants are flowed across the chip in solution. **(b).** Kinetic charts for IL-10Rβ binding for wild type and high affinity IL-10 with inserts for affinity curves. Concentrations used are shown on curves. **(c).** K_D_ values for IL-10Rβ binding for wild type and high affinity variants. **(d).** IL-10Rβ is immobilised on the chip surface and IL-10 variants pre-bound to IL-10Rα are flowed across the chip surface in solution. Concentrations used are shown on curves. **(e).** Kinetics for IL-10Rβ binding in the presence of IL-10Rα. **(f).** K_D_ values for IL-10Rβ binding when IL-10 proteins are pre-bound to IL-10Rα.

**Supplementary Figure 3.**
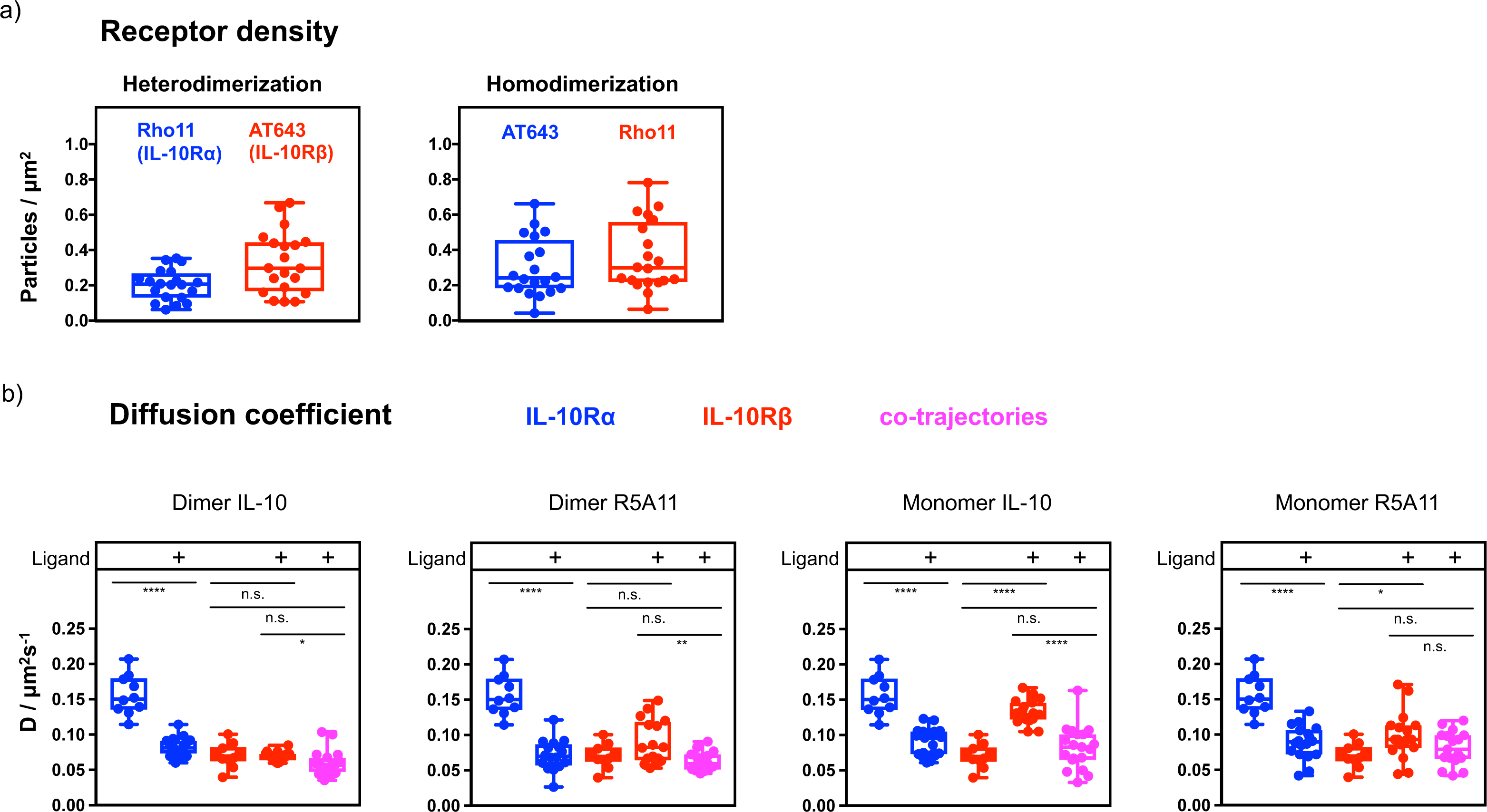
Single molecule imaging of IL-10 receptors by TIRF microscopy: **(a).** Cell surface receptor density of ectopically expressed IL-10Rα (blue) and IL-10Rβ (red). n = 20 cells. **(b).** Diffusion coefficients of IL-10Rα (blue), IL-10Rβ (red) and co-locomoting receptors (magenta) in absence or presence of dimeric and monomeric IL-10 variants. WTD: n = 20 cells; R5A11D: n = 16 cells, WTM: n = 19 cells, R5A11M: n = 18 cells, unstimulated: n = 10 cells.

**Supplementary Figure 4.**
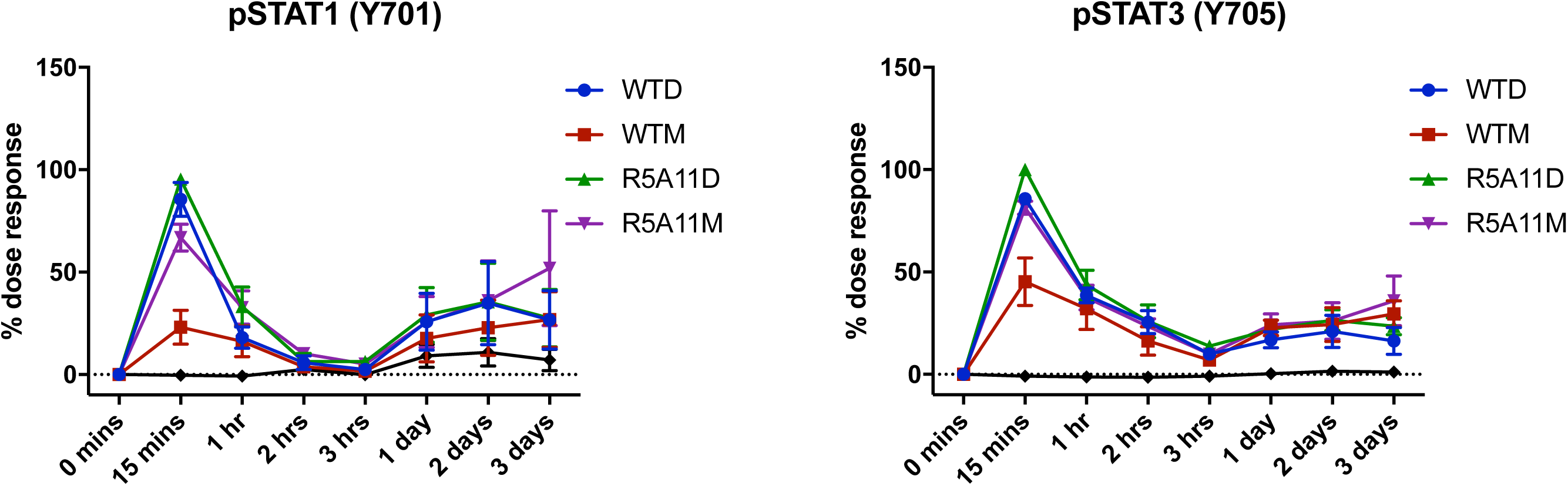
Extended kinetics of IL-10 and variants in human monocytes. 3-day monocyte pSTAT3/1 kinetics. Monocytes were stimulated with IL-10 for the indicated time periods before fixation. Data shown is the mean of four biological replicates with error bars depicting standard error of the mean. Each biological replicate is normalised by assigning the highest MFI value at 15 mins as 100% and the lowest MFI value of an untreated control as 0%.

**Supplementary Figure 5.**
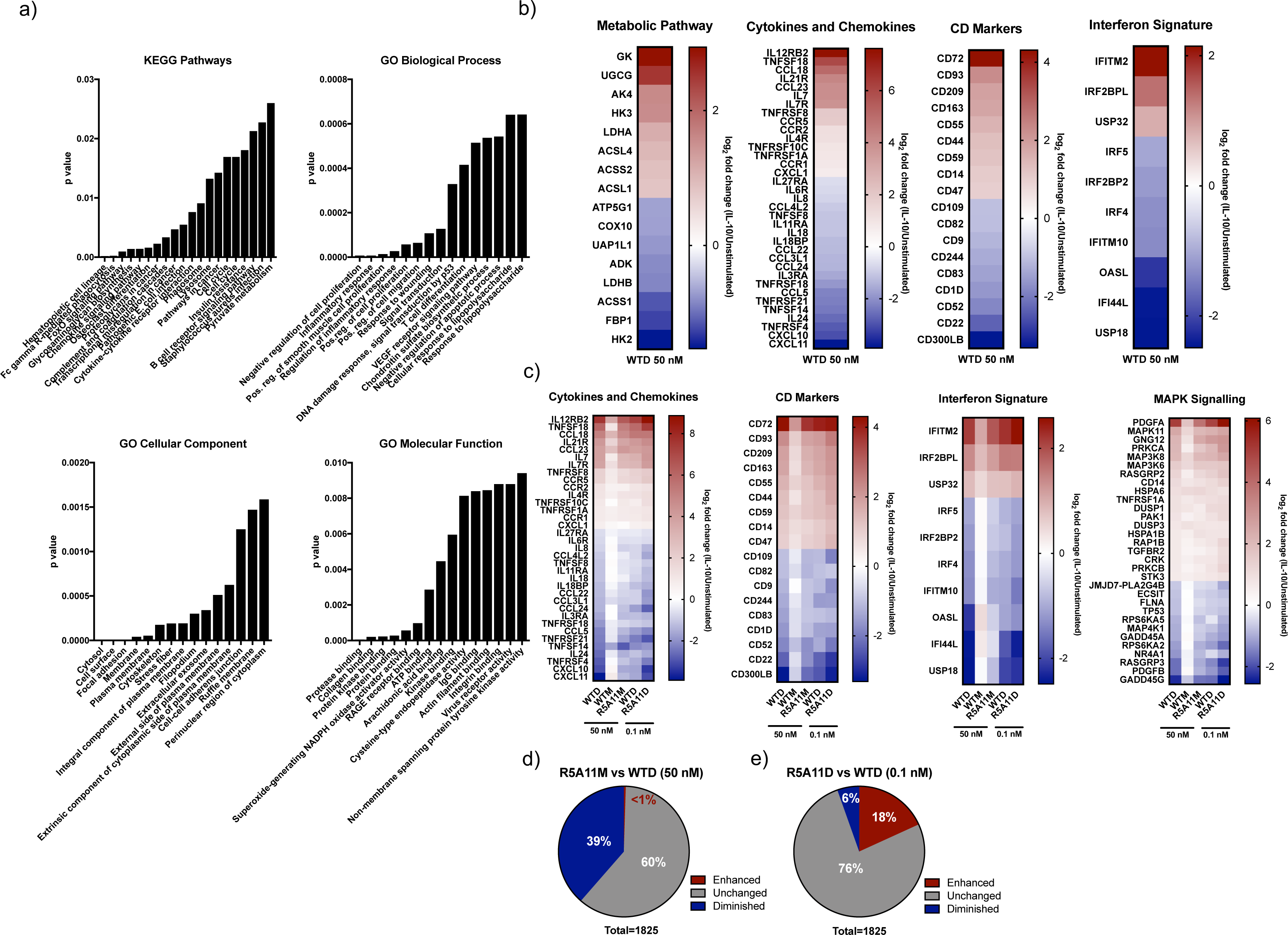
Analysis of gene expression profiles induced by IL-10 wild type and high affinity variants in human monocytes. **(a).** KEGG and GO pathway analysis for genes significantly up or down regulated by WTD 50 nM ≧ 0.6 or ≦ −0.6 log_2_ fold change in human monocytes. Pathway analysis done using DAVID Bioinformatics Resource functional annotation tool (Huang da et al., 2009a, b). **(b).** Heatmap showing log_2_ fold change expression by WTD (50 nM) stimulation for a selection of metabolic pathways, cytokines & chemokines, CD and interferon related genes. **(c).** Comparison of log_2_ fold change in expression for genes by WTD, WTM and R5A11M at 50 nM and WTD and R5A11D at 0.1 nM concentration. **(d).** Comparison of regulation of monocyte genes by R5A11M 50 nM and WTD 50 nM. The log_2_ fold change of R5A11M 50 nM/unstimulated was divided by the log_2_ fold change of WTD 50 nM/unstimulated. Proportion of genes which show enhanced regulation by R5A11M are shown in red, proportion of genes which show diminished regulation by R5A11M are shown in blue and genes which do not change between R5A11M and WTD are shown in grey. **(e).** Comparison of regulation of genes by R5A11D 0.1 nM and WTD 0.1 nM. The log_2_ fold change of R5A11D 0.1nM/unstimulated was divided by the log_2_ fold change of WTD 0.1 nM/unstimulated. Proportion of genes which show enhanced regulation by R5A11D are shown in red, proportion of genes which show diminished regulation by R5A11D are shown in blue and genes which do not change between R5A11D and WTD are shown in grey.

**Supplementary Figure 6.**
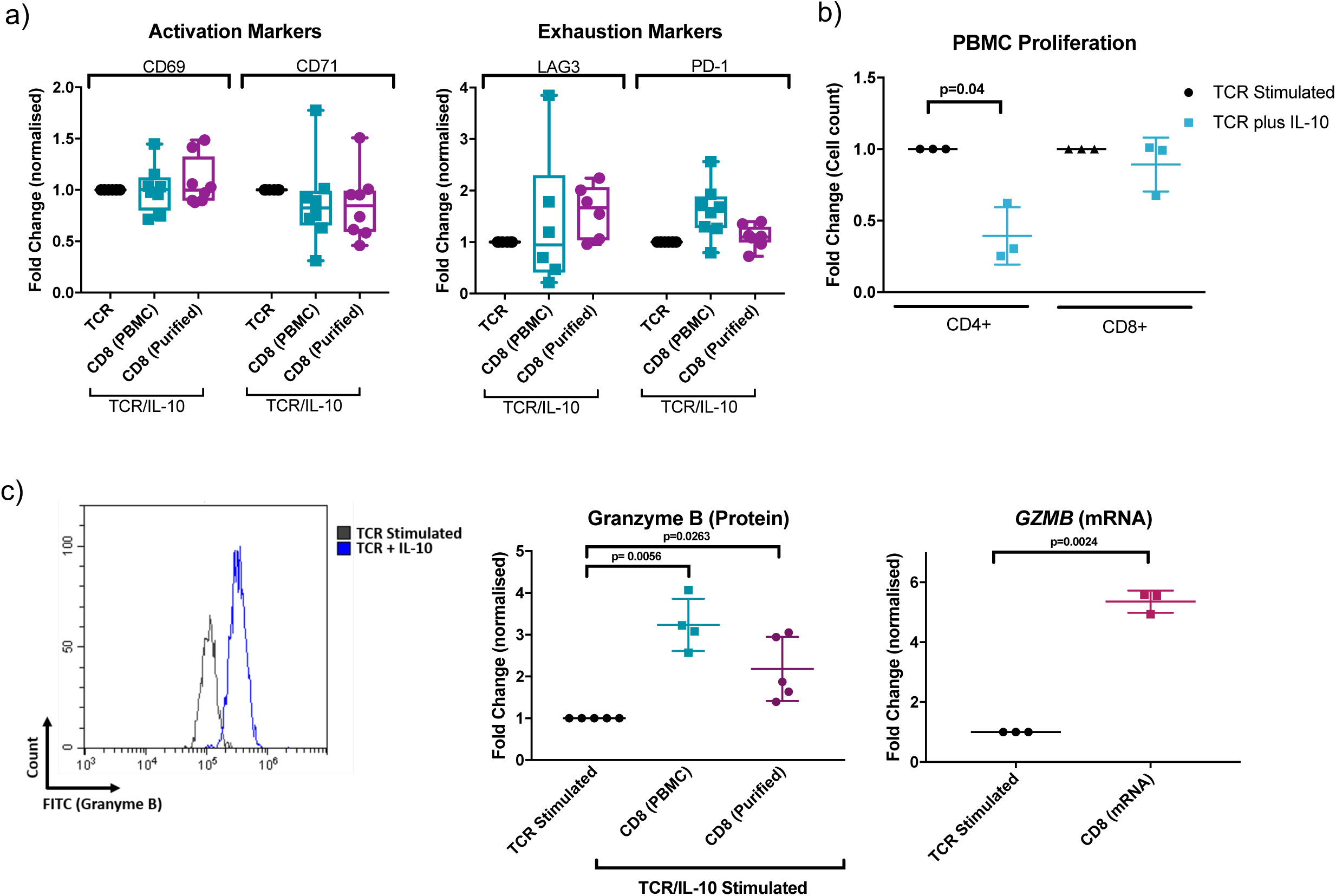
Characterisation of the IL-10 treated CD8 T cell phenotype. **(a).** CD8 cells within a PBMC population and purified CD8 cells were stained for CD69 after 24 hours activation and for CD71 after 6 days activation and expansion. Exhaustion markers PD-1 and LAG3 were analysed after 6 days activation and expansion. Fold change was calculated by dividing IL-10 stimulated MFI values by non-IL-10 stimulated controls for each donor. Each point represents one donor. **(b).** Proliferation of CD4 and CD8 T cells in a PBMC population were analysed after 6 days of activation/expansion. Cell counts for CD4 and CD8 cells were taken and fold change was calculated by dividing IL-10 treated cells by a non-IL-10 treated control population from the same donor. Each point represents one biological replicate and p values were calculated using a two tailed paired t test. **(c).** CD8 T cells in a purified population were stained for granzyme B. Fold change of granzyme B was calculated by normalising within each biological replicate to a non-IL-10 treated control (TCR stimulated) for both CD8 cells in a PBMC population and purified CD8 cells. mRNA was isolated from a purified CD8 cell population and *gzmb* mRNA was quantified by RT qPCR. Fold change was calculated by dividing by a non-IL-10 treated control. Each point represents a biological replicate and p values were calculated using two tailed paired t test.

**Supplementary Figure 7.**
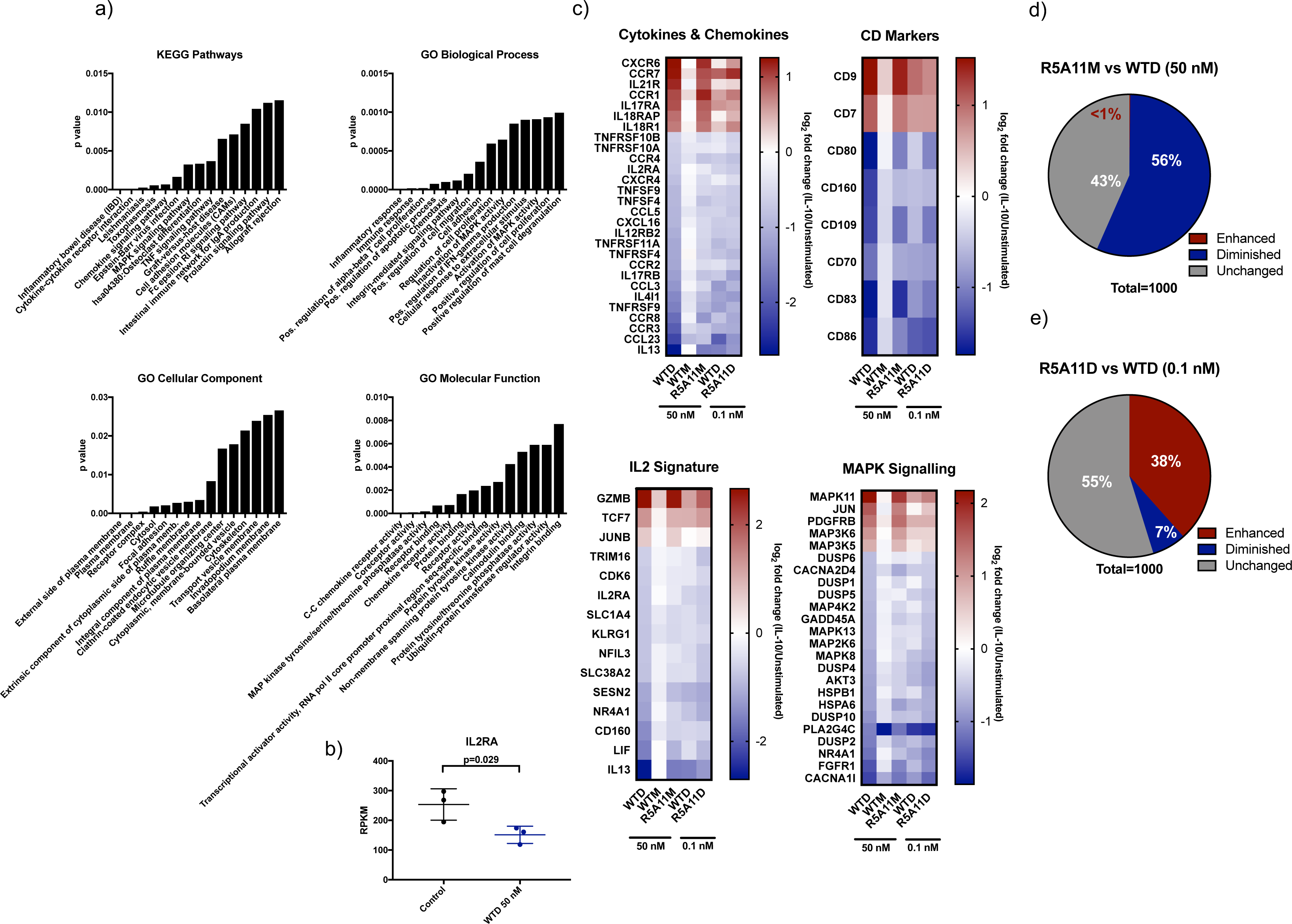
Analysis of gene expression profiles induced by IL-10 wild type and high affinity variants in human CD8 T cells. **(a).** KEGG and GO pathway analysis for genes significantly up or down regulated by WTD 50 nM ≧ 0.6 or ≦ −0.6 log_2_ fold change in human CD8 T cells. Pathway analysis done using DAVID Bioinformatics Resource functional annotation tool (Huang da et al., 2009a, b). **(b).** The RKPM of unstimulated and WTD 50 nM conditions for the IL2RA gene in each donor. **(c).** Heatmap comparison of regulation of cytokines & chemokines, CD markers, IL-2 related and MAPK signaling genes by WTD, WTM and R5A11M at 50 nM and WTD and R5A11D at 0.1 nM. **(d).** The log_2_ fold change of R5A11M 50 nM/unstimulated was divided by the log_2_ fold change of WTD 50 nM/unstimulated. Proportion of genes which show enhanced regulation by R5A11M are shown in red, proportion of genes which show diminished regulation by R5A11M are shown in blue and genes which do not change between R5A11M and WTD are shown in grey. **(e).** Comparison of regulation of genes by R5A11D 0.1 nM and WTD 0.1 nM. The log_2_ fold change of R5A11D 0.1nM/unstimulated was divided by the log_2_ fold change of WTD 0.1 nM/unstimulated. Proportion of genes which show enhanced regulation by R5A11D are shown in red, proportion of genes which show diminished regulation by R5A11D are shown in blue and genes which do not change between R5A11D and WTD are shown in grey.

